# Deep sampling of Hawaiian *Caenorhabditis elegans* reveals high genetic diversity and admixture with global populations

**DOI:** 10.1101/716928

**Authors:** Timothy A. Crombie, Stefan Zdraljevic, Daniel E. Cook, Robyn E. Tanny, Shannon C. Brady, Ye Wang, Kathryn S. Evans, Steffen Hahnel, Daehan Lee, Briana C. Rodriguez, Gaotian Zhang, Joost van der Zwaag, Karin C. Kiontke, Erik C. Andersen

## Abstract

Recent efforts to understand the natural niche of the keystone model organism *Caenorhabditis elegans* have suggested that this species is cosmopolitan and associated with rotting vegetation and fruits. However, most of the strains isolated from nature have low genetic diversity likely because recent chromosome-scale selective sweeps contain alleles that increase fitness in human-associated habitats. Strains from the Hawaii Islands are highly divergent from non-Hawaiian strains. This result suggests that Hawaiian strains might contain ancestral genetic diversity that was purged from most non-Hawaiian strains by the selective sweeps. To characterize the genetic diversity and niche of Hawaiian *C. elegans*, we sampled across the Hawaiian Islands and isolated 100 new *C. elegans* strains. We found that *C. elegans* strains are not associated with any one substrate but are found in cooler climates at high elevations. These Hawaiian strains are highly diverged compared to the rest of the global population. Admixture analysis identified 11 global populations, four of which are from Hawaii. Surprisingly, one of the Hawaiian populations shares recent ancestry with non-Hawaiian populations, including portions of globally swept haplotypes. This discovery provides the first evidence of gene flow between Hawaiian and non-Hawaiian populations. Most importantly, the high levels of diversity observed in Hawaiian strains might represent the complex patterns of ancestral genetic diversity in the *C. elegans* species before human influence.

## Introduction

Over the last 50 years, the nematode *Caenorhabditis elegans* has been central to many important discoveries in the fields of developmental, cellular, and molecular biology. The vast majority of these insights came from the study of a single laboratory-adapted strain collected in Bristol, England known as N2 (Brenner, 1974; Chalfie et al., 1994; Consortium, 1998; Fire et al., 1998; Grishok et al., 2000; Hodgkin and Brenner, 1977; Lee et al., 1993; Sulston et al., 1983). Recent sampling efforts have led to the identification of numerous wild *C. elegans* strains and enabled the study of genetic diversity and ecology of the species (Andersen et al., 2012; Barrière and Félix, 2014; Cook et al., 2016; Félix and Duveau, 2012; Ferrari et al., 2017; Hahnel et al., 2018; Lee et al., 2019; Richaud et al., 2018). The earliest studies of *C. elegans* genetic variation showed that patterns of single-nucleotide variant (SNV) diversity were shared among most wild strains, with the exception of a Hawaiian strain, CB4856, which has distinct and high levels of variation relative to other strains (Koch et al., 2000). Subsequent analyses revealed that *C. elegans* has reduced levels of diversity relative to the obligate outcrossing *Caenorhabditis* species and the facultative selfer *C. briggsae* (Dey et al., 2013; Thomas et al., 2015). The most comprehensive analysis of *C. elegans* genetic diversity to date used data from thousands of genome fragments across a globally distributed collection of 97 genetically distinct strains to show that recent selective sweeps have largely homogenized the genome (Andersen et al., 2012). The authors hypothesized that these selective sweeps might contain alleles that facilitate human-assisted dispersal and/or increase fitness in human-associated habitats. Consistent with the previous analyses, two Hawaiian strains, CB4856 and DL238, did not share patterns of reduced genetic diversity caused by the selective sweeps that affected the rest of the *C. elegans* population – a trend that has held true as the number of Hawaiian strains has increased (Cook et al., 2017, 2016; Hahnel et al., 2018; Lee et al., 2019). Taken together, these studies suggest that the Hawaiian *C. elegan*s population might be more representative of ancestral genetic diversity that existed prior to the selective pressures associated with recent human influence.

To better characterize the genetic diversity of the *C. elegans* species on the Hawaiian Islands, we performed deep sampling across five Hawaiian islands: Kauai, Oahu, Molokai, Maui, and the Big Island. Because incomplete data on locations and environmental parameters are common issues for some field studies of *C. elegans* (Andersen et al., 2012; McGrath et al., 2009; Rockman and Kruglyak, 2009), we developed a standardized collection procedure with the Fulcrum® mobile data collection application. This streamlined procedure enabled us to rapidly record GPS coordinates and environmental niche parameters at each collection site, and accurately link these data with the nematodes we isolated. The Hawaiian Islands are an ideal location to study characteristics of the *C. elegans* niche because the Islands contain many steep, wide-ranging gradients of temperature, humidity, elevation, and landscape usage. In total, we collected samples from 2,263 sites across the islands and isolated 2,532 nematodes, including 309 individuals from the *Caenorhabditis* genus. Among these isolates, we identified 100 new *C. elegans* strains, 95 of which proliferated in the lab and were whole-genome sequenced. Analysis of genomic variation revealed that these strains represent 26 distinct genome-wide haplotypes not sampled previously. We refer to these genome-wide haplotypes as isotypes. We grouped these 26 Hawaiian isotypes with the 17 previously isolated Hawaiian isotypes and compared their genetic variation to 233 non-Hawaiian isotypes from around the globe. Consistent with previous observations, we found that the Hawaiian population has approximately three times more diversity than the non-Hawaiian population. However, we were surprised to find that, in a subset of Hawaiian isotypes, some genomic regions appear to be shared with non-Hawaiian isotypes from around the globe. These results provide the first evidence of gene flow between these populations and suggest that future sampling efforts in the Hawaiian Islands will help elucidate the evolutionary processes that have shaped the genetic diversity in the *C. elegans* species.

## Results

### Hawaiian nematode diversity

In August 2017, we collected a total of 2,263 samples across five Hawaiian islands and ascertained the presence of nematodes in each sample (**Figure 1, Supplemental Table 5**). We isolated one or more nematodes from 1,120 of 2,263 (48%) samples, and an additional 431 of 2,263 (19%) samples had circumstantial evidence of nematodes (tracks but no nematodes could be found on the collection plate). Altogether, we isolated 2,531 nematodes from 1,120 samples and genotyped them by analysis of the Internal Transcribed Spacer (ITS2) region between the 5.8S and 28S rDNA genes (Barrière and Félix, 2014; Kiontke et al., 2011). We refer to isolates where the ITS2 region was amplified by PCR as ‘PCR-positive’ and isolates with no amplification as ‘PCR-negative’ (see Methods). The PCR-positive category comprises *Caenorhabditis* isolates that we identified to the species level and isolates from genera other than *Caenorhabditis* that we identified to the genus level. Using this categorization strategy, we found that 427 of 2,531 isolates (17%) were PCR-positive and belonged to 13 distinct taxa. Among all isolates, we identified five *Caenorhabditis* species at different frequencies across the 2,263 samples: *C. briggsae* (4.2%), *C. elegans* (1.7%), *C. tropicalis* (0.57%), *C. kamaaina* (0.088%), and a new species *C. oiwi* (0.53%) (**Supplemental Table 5**). We named *Caenorhabditis oiwi* for the Hawaiian word meaning “native” in reference to its endemic status on the Hawaiian Islands. This species was found to be distinct based on molecular barcodes (Kiontke et al., 2011) and on biological species inference from mating crosses (Félix et al., 2014) (**Supplemental File 1**). The most common *Caenorhabditis* species we isolated was *C. briggsae*, which is consistent with nematode collection efforts by other groups that suggest *C. briggsae* is a ubiquitous species in many regions of the world (Félix et al., 2013). We found no evidence of island enrichment for *Caenorhabditis* species apart from *C. elegans*, where it was enriched on the Big Island relative to Kauai and Maui (Fisher’s Exact Test, *p* < 0.01).

**Figure 1.**
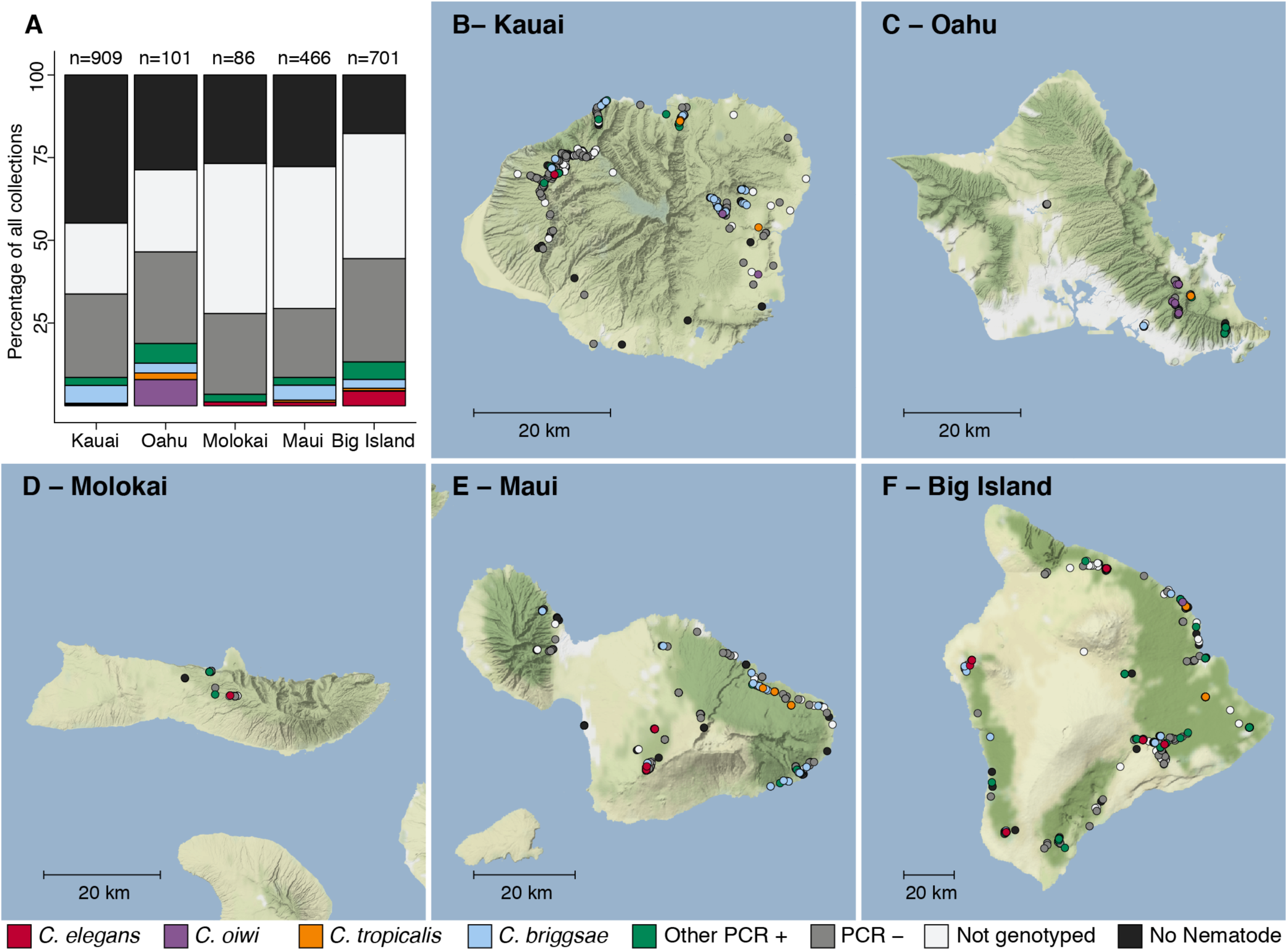
Geographic distribution of sampling sites across five Hawaiian islands. In total we sampled 2,263 unique sampling sites. (**A**) The percentage of each collection category is shown by island. The collection categories are colored according to the legend at the bottom of the panel, and the total number of samples for each island are shown above the bars. (**B-F**) The circles indicate unique sampling sites (n = 2,263) and are colored by the collection categories shown in the bottom legend. For sampling sites where multiple collection categories apply (n = 299), the site is colored by the collection category shown in the legend from left to right, respectively. For all sampling sites, the GPS coordinates and collection categories found at that site are included in (**Supplemental Data 1**). We focused our studies on *Caenorhabditis* nematode collections, excluding *C. kamaania* because it was only found at two sampling sites. Maps © www.thunderforest.com, Data © www.osm.org/copyright.

### The *C. elegans* niche is distinct from other *Caenorhabditis* species on Hawaii

To characterize more about a nematode niche on the Hawaiian Islands, we classified the substrate for each distinct collection and measured various environmental parameters. Of the six major classes of substrate, we found nematodes most often on leaf litter (56%). When we account for collections with nematode-like tracks on the collection plate, we estimated that greater than 80% of leaf litter substrates contained nematodes (**Figure 2A**). The isolation success rate for the other classes of substrate ranged from 35% to 48% (**Figure 2A**). In comparison to overall nematode isolation rates, *Caenorhabditis* nematodes were isolated more frequently from flower substrates (40 of 202 collections) than any other substrate category (Fisher’s Exact Test, *p* < 0.02) (**Figure 2A**). We also found that *Caenorhabditis* nematodes were enriched on rotting fruits, nuts, or vegetables (33 of 333 collections) relative to leaf litter substrates (76 of 1480 collections) (Fisher’s Exact Test, *p* < 0.02) but not other substrate classes (**Figure 2A**). These findings are consistent with other collection surveys that have shown leaf litter substrates harbor fewer *Caenorhabditis* nematodes than rotting flowers and fruits (Félix et al., 2013; Ferrari et al., 2017). We observed similar trends of flower-substrate enrichment relative to leaf litter for *C. briggsae* (Fisher’s Exact Test, *p* = 0.00049; flower, 21 of 202 collections and leaf litter 51 of 1480 collections) and *C. tropicalis* (Fisher’s Exact Test, *p* = 0.0059; flower, five of 202 collections and leaf litter, three of 1480 collections) but not for *C. elegans* (Fisher’s Exact Test, *p* = 1), which exhibited no substrate enrichment (**Figure 2B-C**). Interestingly, the new species, *C. oiwi*, was only isolated from flower and fruit/nut/vegetable substrates and was enriched on flower substrates (Fisher’s Exact Test, *p* = 0.0124; flower, nine of 202 collections and fruit/nut/vegetable, three of 333 collections) (**Figure 2B-C**).

**Figure 2.**
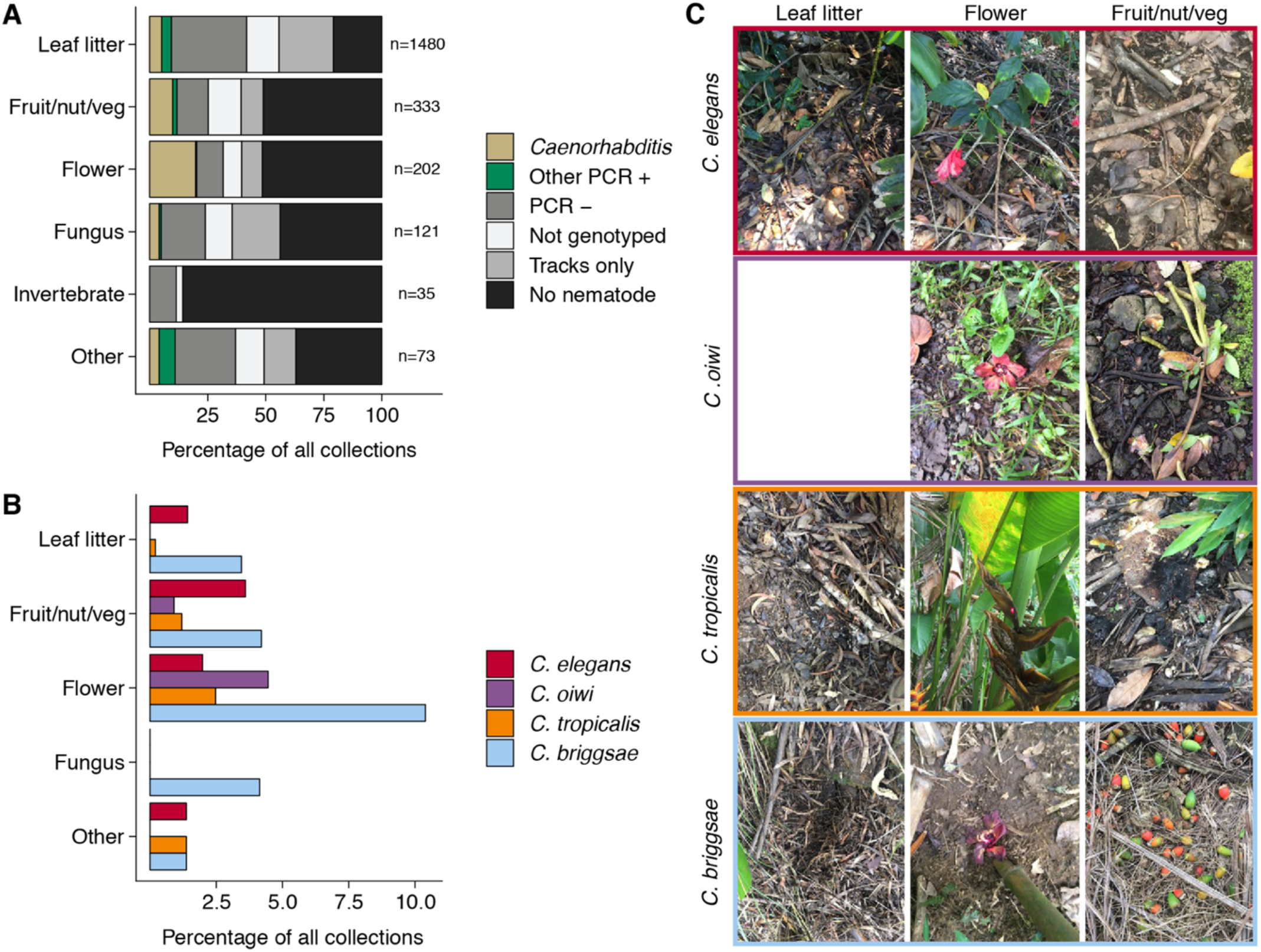
Collection categories by substrate type. (**A**) The percentage of each collection category is shown by substrate type. The collection categories are colored according to the legend at the right, and the total number of samples for each substrate are shown to the right of bars. (**B**) The percentage of collections is shown by substrate type for each *Caenorhabditis* species (excluding *C. kamaaina*, n = 2). (**C**) Examples of substrate photographs for *Caenorhabditis* species are shown. The *C. oiwi* leaf litter cell is blank because *C. oiwi* was only isolated from flowers and fruit.

The enrichment of *C. briggsae, C. tropicalis*, and *C. oiwi* on flowers might indicate that this substrate class has a higher nutrient quality for these species. If this hypothesis is correct, we might expect to see a greater incidence of proliferating populations on flower substrates than other substrates. However, we saw no observable association between large population size (approximate number of nematodes on collection plate) and substrate class for *C. briggsae* (Spearman’s *rho* = −0.0197, *p* = 0.57 flower vs. leaf litter), *C. tropicalis* (Spearman’s *rho* = −0.26, *p* = 0.73 flower vs. leaf litter), nor *C. oiwi* (Spearman’s *rho* = 0.258, *p* = 0.21 flower vs. fruit/nut/vegetable), which suggests that other factors might drive the observed flower enrichment or that we are limited by the small sample size. Taken together, these data suggest that the *Caenorhabditis* species we isolated do not exhibit substrate specificity, despite flower-substrate preferences of *C. briggsae, C. tropicalis*, and *C. oiwi*, which is different from some other species in the genus that demonstrate substrate specificity (*e.g., C. astrocarya* and *C. inopinata*) (Ferrari et al., 2017; Kanzaki et al., 2018).

In addition to recording substrate classes, we measured elevation, ambient temperature and humidity, and substrate temperature and moisture to determine if these niche parameters were important for individual *Caenorhabditis* species (**Figure 3**; see Methods).

**Figure 3.**
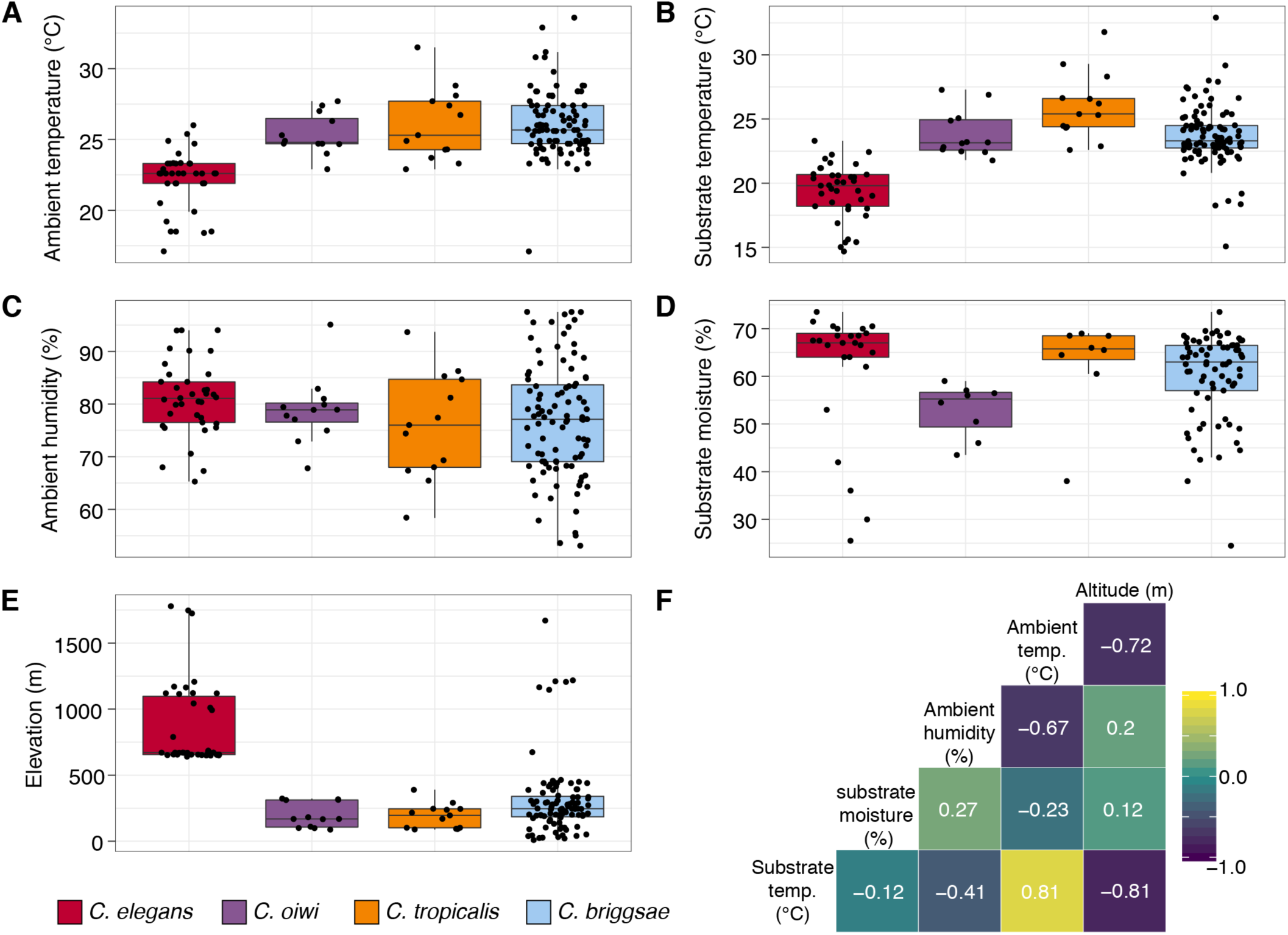
Environmental parameter values for sites where *Caenorhabditis* species were isolated. (**A-E**) Tukey box plots are plotted by species (colors) for different environmental parameters. Each dot corresponds to a unique sampling site where that species was identified. In cases where two *Caenorhabditis* species were identified from the same sample (n = 3), the same parameter values are plotted for both species. All p-values were calculated using Kuskal-Wallis test and Dunn test for multiple comparisons with *p* values adjusted using the Bonferroni method; comparisons not mentioned were not significant (α = 0.05). (**A**) Ambient temperature (°C) was typically cooler at the sites were *C. elegans* were isolated compared to sites for all other *Caenorhabditis* species (Dunn test, *p* < 0.005). (**B**) Substrate temperature (°C) was also generally cooler for *C. elegans* than all other *Caenorhabditis* species (Dunn test, *p* < 0.00001). (**C**) Ambient humidity (%) did not differ significantly among the *Caenorhabditis-*positive sites. (**D**) Substrate moisture (%) was generally greater for *C. elegans* than *C. oiwi* (Dunn test, *p* = 0.002). (**E**) Elevation (meters) was typically greater at sites where *C. elegans* were isolated compared to sites for all other *Caenorhabditis* species (Dunn test, *p* < 0.00001). (**F**) A correlation matrix for the environmental parameters was made using sample data from the *Caenorhabditis* species shown. The parameter labels for the matrix are printed on the diagonal, and the pearson correlation coefficients are printed in the cells. The color scale also indicates the strength and sign of the correlations shown in the matrix.

Consistent with previous *C. elegans* collections in tropical regions (Andersen et al., 2012; Dolgin et al., 2008), all *C. elegans* isolates were collected from elevations greater than 500 meters and were generally found at higher elevations than other *Caenorhabditis* species (**Figure 3E;** mean = 867 m; elevation: Dunn test, *p* < 0.00001). We also found that *C. elegans-*positive collections tended to be at cooler ambient and substrate temperatures than other *Caenorhabditis* species (ambient temperature: Dunn test, *p* < 0.005; substrate temperature: Dunn test, *p* < 0.00001), although these two environmental parameters were correlated with elevation (**Figure 3F**). Notably, the average substrate temperatures for *C. elegans* (19.4 °C), *C. tropicalis* (26.0 °C), and *C. briggsae* (23.7°C) positive collections are close to the optimal growth temperatures for these species in the laboratory setting (**Figure 3B**) (Poullet et al., 2015). Our collections also indicate that *C. oiwi* tends to be found on drier substrates than *C. elegans* (**Figure 3D**; Dunn test, *p* = 0.0021), but we observed no differences among species for ambient humidity (**Figure 3C**). Given the similar substrate and environmental parameter preferences of *C. tropicalis, C. briggsae*, and *C. oiwi*, we next asked if these species colocalized at either the local (< 30 m^2^) or substrate (< 10 cm^2^) scales. To sample at the local scale, we collected samples from 20 gridsects (see Methods; **Supplemental Figure 1**) and observed no colocalization of these three species, although only 16% of the total collections were a part of a gridsect. At the substrate scale, we found *C. tropicalis* and *C. briggsae* cohabitating on two of 108 substrates with either species present and *C. oiwi* and *C. briggsae* cohabitating on one of 107 substrates with either species present (**Supplemental Figure 2**). Among 95 substrates with *C. briggsae*, we observed nine instances of *C. briggsae* cohabitating with other PCR-positive species. We did not collect any samples that harbored *C. elegans* and any other *Caenorhabditis* species. Taken together, these cohabitation results highlight the ubiquitous nature of *C. briggsae* on the Hawaiian Islands and further suggests that the niche of *C. elegans* might be distinct from *C. tropicalis, C. briggsae*, and *C. oiwi* on the Hawaiian Islands.

### Hawaiian *C. elegans* are divergent from the global population

We previously showed that two *C. elegans* isolates from Hawaii are highly divergent relative to wild isolates from other regions of the world and represent a large portion of the genetic diversity found within the species (Andersen et al., 2012). Since this analysis, an additional 15 isolates have been collected from the islands and show similarly high levels of genetic diversity (Cook et al., 2016; Hahnel et al., 2018). To better characterize the genetic diversity in Hawaii, we acquired whole-genome sequence data from 95 *C. elegans* isolates that we collected in this study. By analyzing the variant composition of these 95 isolates, we identified 26 distinct genome-wide haplotypes that we refer to as isotypes (see Methods). Within these 26 isotypes, we identified approximately 1.54 million single nucleotide variants (SNVs) that passed our filtering strategy (see Methods; hard-filter VCF**; Supplemental Table 4**), which is 27.6% greater than the total number of SNVs identified in all of the 233 non-Hawaiian isotypes included in this study. We found that distinct isotypes are frequently isolated within close proximity to one another in Hawaii. We identified up to seven unique isotypes colocalized within a single grisect (less than 30 m^2^) (**Supplemental Figure 3A**). We also found that colocalization occurred at the substrate level; among the 38 substrates from which we isolated *C. elegans*, 12 contained two or more isotypes (**Supplemental Figure 3B**). The variant data from all 43 Hawaiian isotypes (26 new with 17 previously described Hawaiian isotypes) allowed us to perform detailed analyses of Hawaiian genetic diversity.

Consistent with what is known about the *C. elegans* global population (Andersen et al., 2012), we observed a high degree of genome-wide relatedness among a majority of non-Hawaiian isotypes (**Supplemental Figure 4**). By contrast, the Hawaiian isotypes are all diverged from the non-Hawaiian population with the exception of five non-Hawaiian isotypes. Among these exceptions, ECA36 and QX1211 were collected from urban gardens in New Zealand and San Francisco, CA respectively, and grouped with some of the most divergent isotypes from Hawaii. More surprisingly, three non-Pacific Rim isotypes also grouped with the Hawaiian isotypes. These include JU2879, MY16, and MY23. JU2879 was isolated from a rotting apple in Mexico City, Mexico and both MY isotypes were isolated from garden composts in Nordrhein-Westfalen, Germany, separated by approximately 5 km. Within the Hawaiian population, genome-wide relatedness revealed a high degree of divergence (**Supplemental Figure 4**). This trend is further supported by elevated levels of genome-wide average nucleotide diversity (π) in the Hawaiian population relative to the non-Hawaiian population, which we found to be three-fold higher (Hawaii π = 0.00124; non-Hawaiian π = 0.000408, **Figure 4A; Supplemental Data 2)**.

The genomic distribution of diversity followed a similar pattern across chromosomes for both populations, wherein chromosome centers and tips exhibited lower diversity on average than chromosome arms (**Figure 4A; Supplemental Data 2**). This pattern is likely explained by lower recombination rates, higher gene densities, and elevated levels of background selection on chromosome centers (Consortium, 1998; Cutter and Payseur, 2003; Rockman et al., 2010). Interestingly, we observed discrete peaks of diversity in specific genomic regions (*e.g.*, chr IV center), which suggests that balancing selection might maintain diversity at these loci in both populations (**Figure 4A; Supplemental Data 2**). This hypothesis is supported by corresponding spikes in Tajima’s *D* (**Figure 4B; Supplemental Data 3**) (Tajima, 1989). Alternatively, higher values of Tajima’s *D* might indicate a population contraction, but the discrete nature of these peaks makes this possibility less likely. A third possible explanation is that uncharacterized structural variation (*e.g.*, duplication and divergence) exists in these regions. Nevertheless, the variant sites within these discrete peaks in π and Tajima’s *D* are unlikely the result of sequencing errors because they are identified across multiple samples (see Methods).

**Figure 4.**
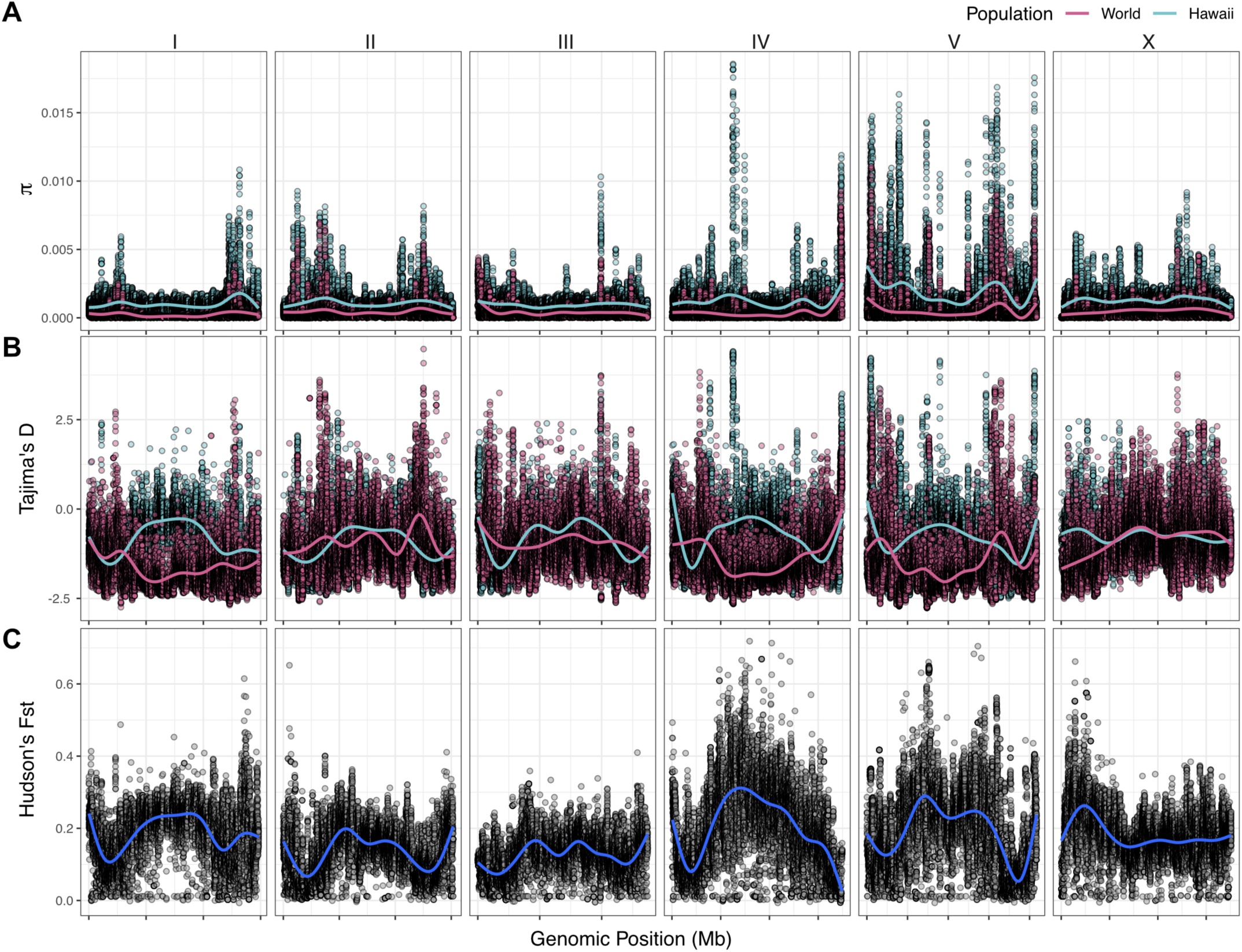
Chromosomal patterns of *C. elegans* diversity and divergence. All comparisons are between the 43 Hawaiian isotypes and the 233 isotypes from the rest of the world. All statistics were calculated along a sliding window of size 10 kb with a step size of 1 kb. Each dot corresponds to the calculated value for window. (**A**) Genome-wide π calculated for Hawaiian isotypes (light blue) and non-Hawaiian isotypes (pink) are shown. (**B**) Genome-wide Tajima’s *D* statistics for Hawaiian isotypes (light blue) and non-Hawaiian isotypes (pink) are shown. (**C**) Genome-wide Hudson’s *F*_*ST*_ comparing the Hawaiian and non-Hawaiian isotypes are shown.

Our previous analysis showed that 70–90% of isotypes contain reduced levels of diversity across several megabases (Mb) on chromosomes I, IV, V, and X (Andersen et al., 2012). This reduced diversity was hypothesized to be caused by selective sweeps that occurred within the last few hundred years, potentially through drastic alterations of global environments by humans. The two Hawaiian isotypes, CB4856 and DL238, did not share this pattern of reduced diversity, suggesting that they avoided the selective pressure. Consistent with this previous analysis, we did not observe signatures of selection in the Hawaiian population on chromosomes I, IV, V, and X, as measured by Tajima’s *D* (**Figure 4B; Supplemental Data 3**), which suggests that the Hawaiian and non-Hawaiian populations have distinct evolutionary histories. This distinction is also captured in genome-wide Hudson’s *F*_*ST*_, where the divergence between the two populations is highest in regions of the genome impacted by the selective sweeps (**Figure 4C; Supplemental Data 2**) (Bhatia et al., 2013; Hudson et al., 1992). Taken together, these data suggest that the Hawaiian population has largely been isolated from the selective pressures thought to be associated with human activity in many regions of the world.

### *C. elegans* population structure on Hawaii

To assess population structure among all 276 isotypes, we performed admixture analysis (see Methods). This analysis suggested that the *C. elegans* species is composed of at least 11 ancestral populations (K), as indicated by the minimization of cross-validation (CV) error between Ks 11-15 (**Supplemental Figure 5**). The population assignments for K=11 closely aligned to the relatedness clusters we observed in a neighbor-joining network of all Hawaiian strains and the species-wide tree (**Figure 5, Supplemental Figure 4**). For Ks 11-15, the majority of Hawaiian isotypes consistently exhibit no admixture with non-Hawaiian ancestral populations. However, a minority of Hawaiian isotypes are consistently either admixed with non-Hawaiian populations (e.g. K=11, 14, and 15) or assigned to ancestral populations that contain non-Hawaiian isotypes (e.g. K=12 and 13) (**Supplemental Figure 5**). These data support that a subset of Hawaiian isotypes are consistently shown to exhibit a greater degree of genetic relatedness with non-Hawaiian isotypes across different population subdivisions. Together, we found at least four distinct subpopulations on the Hawaiian Islands and at least seven additional non-Hawaiian subpopulations comprise the remainder of subpopulations from around the globe (**Supplemental Figure 6**).

**Figure 5.**
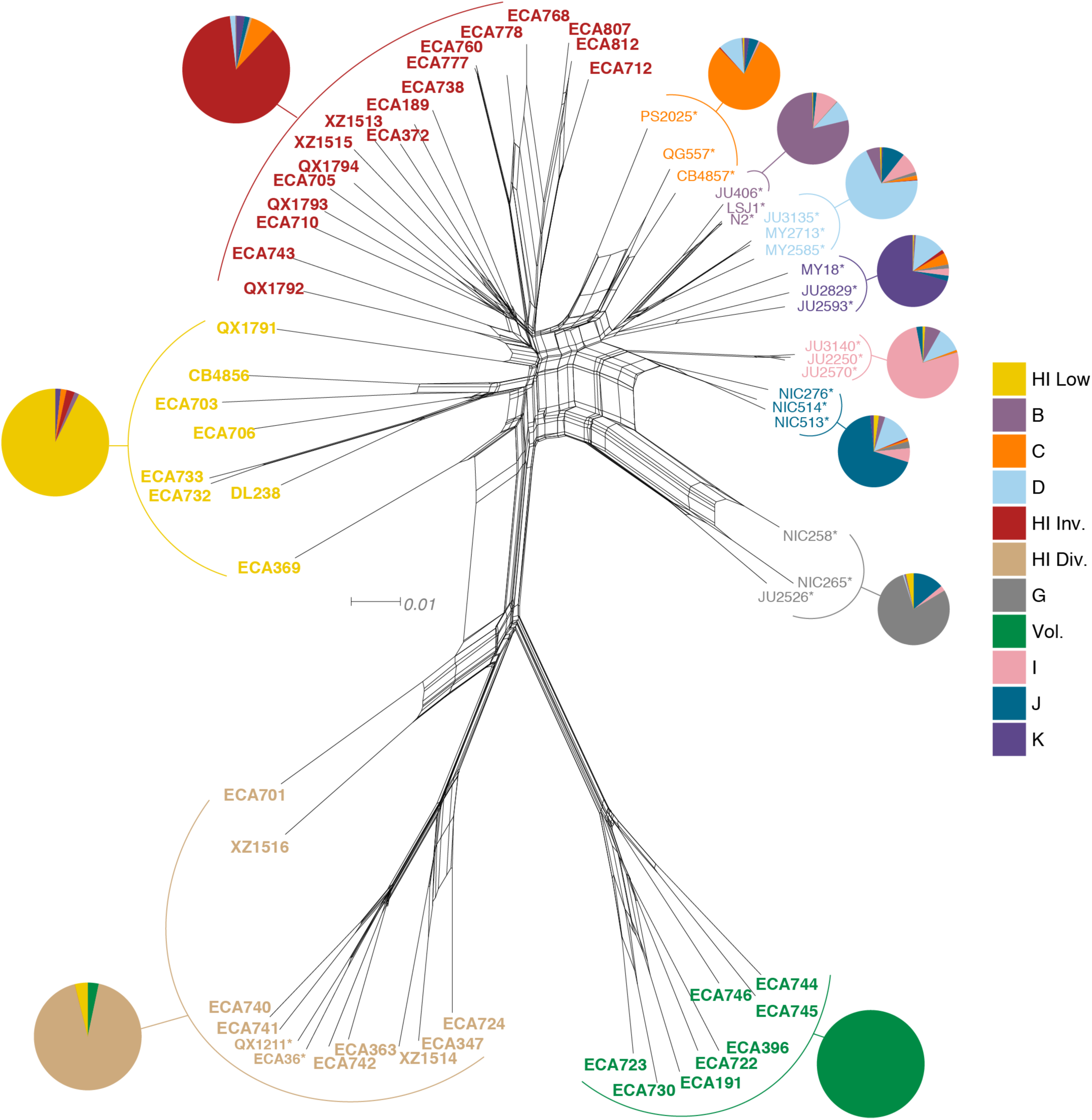
Relatedness of the Hawaiian *C. elegans* isotypes. Neighbor-joining net showing the genetic relatedness of the Hawaiian *C. elegans* population relative to a representative set of non-admixed, non-Hawaiian individuals from each population defined by ADMIXTURE (K=11). Colors of labels indicate the ancestral population assignment from ADMIXTURE (K=11), including the seven global populations (B-K) and the four Hawaiian populations: Hawaiian Invaded, Hawaiian Low, Hawaiian Divergent, and Volcano. Isotypes labeled with an asterisk are representative of non-admixed, non-Hawaiian isotypes from each population defined by ADMIXTURE (K=11). Pie charts represent ancestral population proportions for all isotypes within the full admixture population.

**Figure 6.**
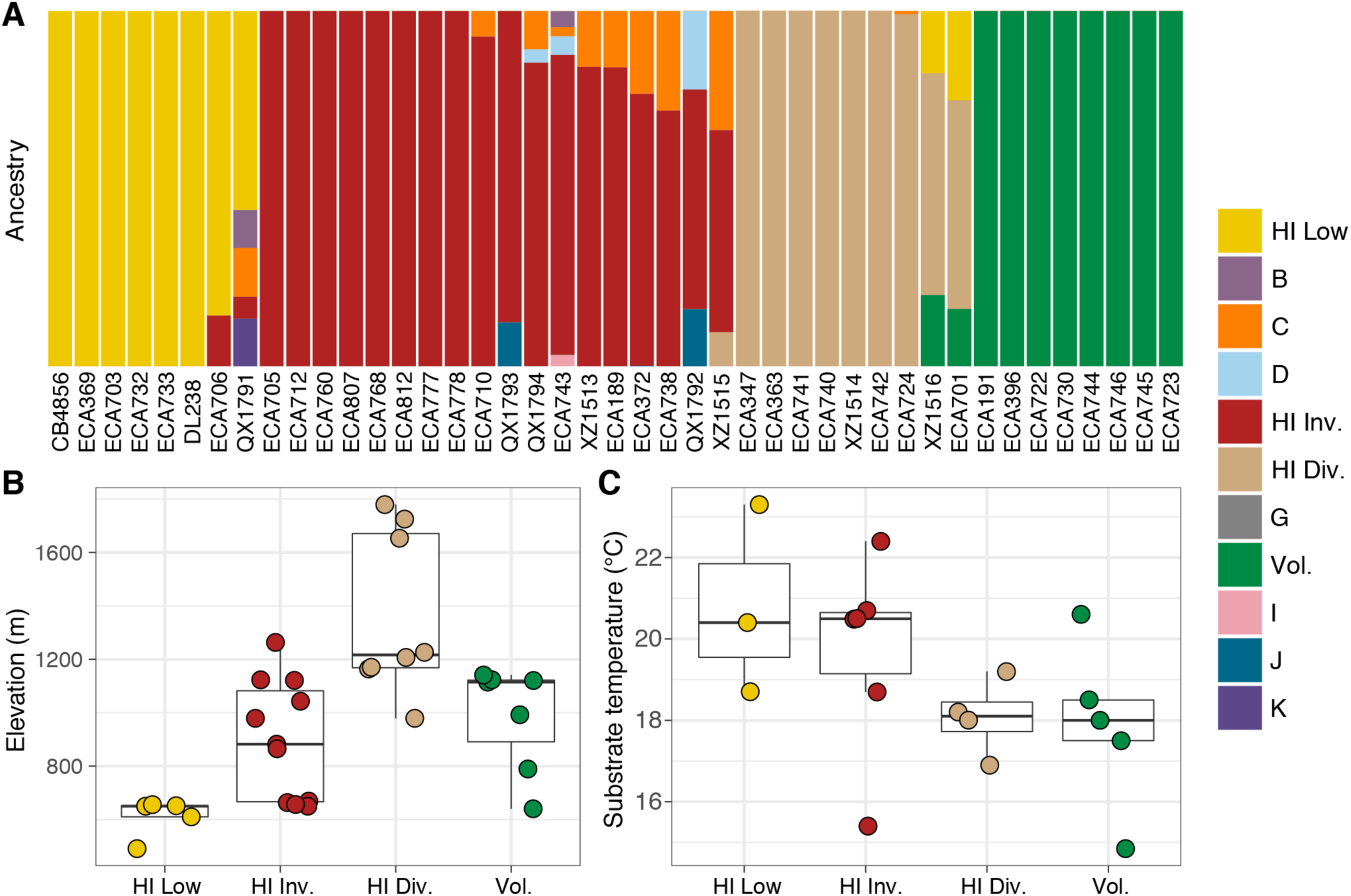
Environmental parameters of Hawaiian *C. elegans* isotypes. (**A**) The inferred ancestral population fractions for each Hawaiian isotype as estimated by ADMIXTURE (K=11; run on the entire *C. elegans* population) are shown. The bar colors represent the ADMIXTURE population assignment for the isotypes named on the x-axis. (**B-C**) Tukey box plots are shown by ADMIXTURE population assignments (colors) for different environmental parameters. We used the average values of environmental parameters from geographically clustered collections to avoid biasing our results by local oversampling (See Methods -Environmental parameter analysis). All p-values were calculated using Kuskal-Wallis test and Dunn test for multiple comparisons with *p* values adjusted using the Bonferroni method; comparisons not mentioned were not significant (α = 0.05). (**B**) The collection site elevations for Hawaiian isotypes colored by the ADMIXTURE population assignments are shown. The Hawaiian Low and the Hawaiian Invaded populations were typically found at lower elevations than the Hiawaiian Divergent population (Dunn test, *p*-values = 0.000168, and 0.037 respectively). (**C**) The substrate temperatures for Hawaiian isotypes colored by the ADMIXTURE population assignments are shown.

The majority of isotypes assigned to the seven non-Hawaiian ancestral populations exhibit a high degree of admixture with one another (at K=11), indicating that these populations are not well differentiated. By contrast, isotypes assigned to three of the four Hawaiian ancestral populations showed almost no admixture. We refer to the four Hawaiian populations as Volcano, Hawaiian Divergent, Hawaiian Invaded, and Hawaiian Low for the following reasons. All eight isotypes in the Volcano population were isolated on the Big Island of Hawaii at high elevation in wet rainforests primarily composed of ferns, ‘Ōhi‘a lehua, and koa trees. We chose to name this population ‘Volcano’ because the majority of isotypes were isolated from the town of Volcano. The Hawaiian Divergent population is named for the two highly divergent isotypes, XZ1516 and ECA701, which were isolated from Kauai, the oldest Hawaiian island sampled. However, we emphasize that the population assignment of these two highly divergent isotypes might not be correct given that they each contain many unique variants that were filtered from the admixture analysis. The Hawaiian Invaded population is named because many of the isotypes assigned to this population exhibited admixture with non-Hawaiian ancestral populations, which is suggestive of an invasion of non-Hawaiian alleles into Hawaii (**Figure 6A, Supplemental Figure 7**).

**Figure 7.**
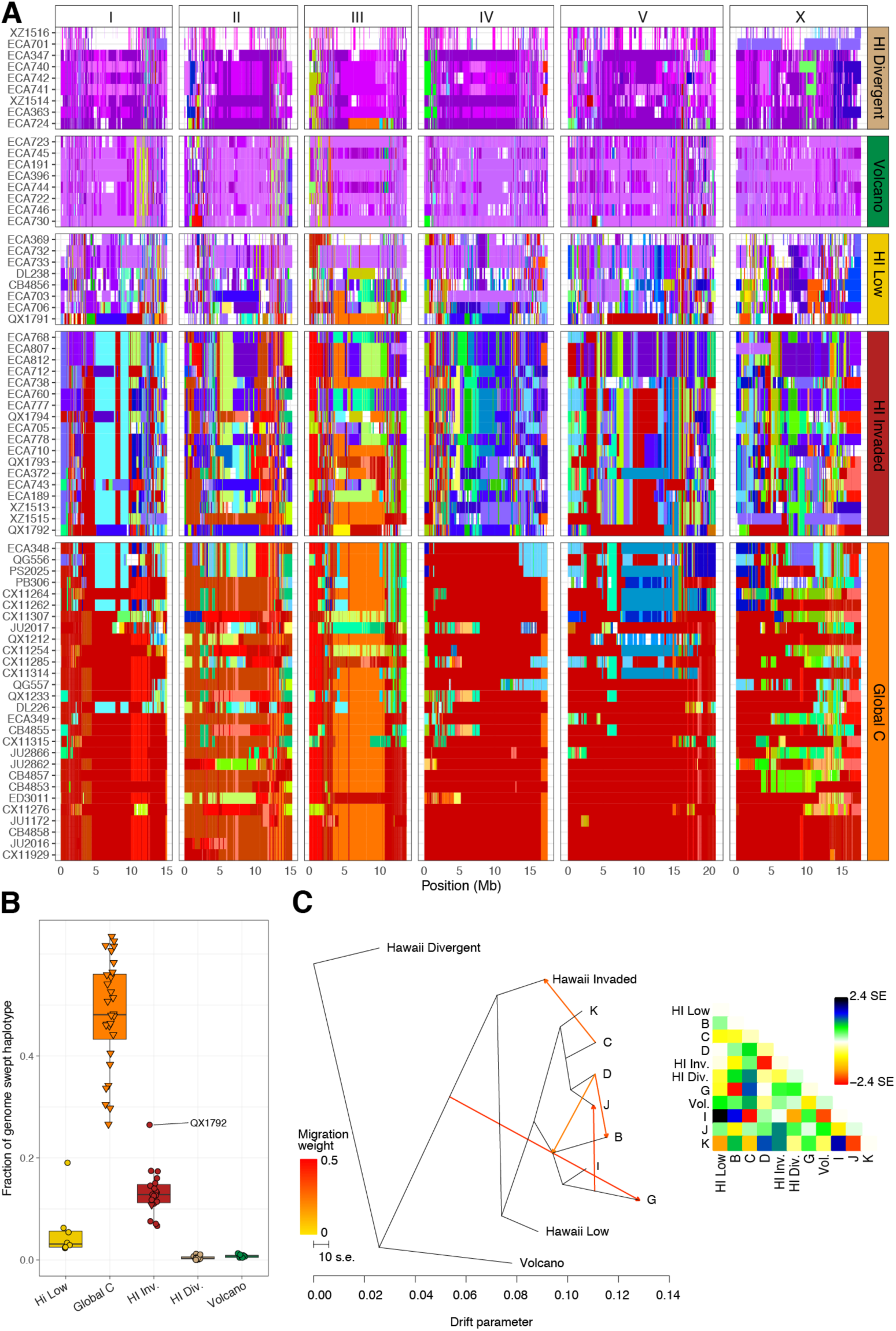
Evidence of migration between the Hawaiian and world populations. (**A**) The inferred blocks of identity by descent (IBD) across the genome are shown. The genomic position is plotted on the x-axis for each isotype plotted on the y-axis. The block colors correspond to a uniquely defined IBD group. The dark red blocks correspond to the most common global haplotype (*i.e.*, the swept haplotype on chr I, IV, V, and left of X). Genomic regions with no color represent regions for which no IBD groups could be determined. The four Hawaiian populations are shown in the top four facets excluding non-Hawaiian isotypes. The bottom facet shows the global C population. (**B**) The total fraction of the genome with the swept haplotype is shown by ancestral population. The data points correspond to isotypes and are colored by their assigned ancestral populations. The Hawaiian isotypes are plotted as circles and non-Hawaiian isotypes are plotted as triangles. Hawaiian isotypes with greater than 25% of their genome swept are labelled. (**C**) The inferred relationship among the ancestral populations allowing for five migration events (ADMIXTURE, K=11). The heat map to the right represents the residual fit to the migration model.

The Hawaiian Low population is named because isotypes assigned to this population tended to be isolated at lower elevations than those assigned to the other Hawaiian populations (See Methods, **Figure 6B**). The population structure of the Hawaiian isotypes suggests that geographic associations within the Hawaiian *C. elegans* population exist either by elevation or by island.

Within the Hawaiian Invaded population, one of the 19 isotypes was isolated from outside of Hawaii (MY23), and 11 of 18 Hawaiian isotypes showed admixture with various non-Hawaiian populations, particularly the non-Hawaiian population C (**Figure 6, Supplemental Figure 6**). By contrast, just one individual assigned to the global C population was admixed with the Hawaiian Invaded population (**Supplemental Figure 6**). This result suggested that these populations either share ancestry or recent gene flow occurred between them. To distinguish between these possibilities, we explicitly tested for the presence of gene flow among all subpopulations using TreeMix (Pickrell and Pritchard, 2012), which estimates the historical relationships among populations accounting for both population splits and migration events. We found evidence of gene flow between the Hawaiian Invaded population and the non-Hawaiian population C (**Figure 7C; Supplemental Figure 8**). The topological position of the fourth highest-weight migration event identified by TreeMix (*i.e.*, C→Hawaiian Invaded) suggested that the evidence of gene flow is not caused by incomplete assortment of ancestral alleles (*i.e.*, the migration arrows connect the ‘C’ and Hawaiian Invaded linages at the branch tips) (**Figure 7C; Supplemental Figure 8**). Importantly, TreeMix cannot distinguish the direction of migration between these subpopulations.

To further assess evidence of gene flow between the Hawaiian and globally distributed subpopulations, we analyzed the haplotype structure across the genomes of all 276 *C. elegans* isotypes (Browning and Browning, 2016). Within the Hawaiian Divergent, Volcano, and Hawaiian Low populations, we observed haplotypes that were largely absent from the non-Hawaiian isotypes. By contrast, the Hawaiian Invaded population shared haplotypes that were commonly found in non-Hawaiian isotypes assigned to non-Hawaiian populations. For example, the isotypes in the Hawaiian Invaded population exhibiting admixture with the global C population share haplotype arrangements on the left and center of Chr III (red and orange Chr III, **Figure 7A**). We also found evidence of the globally swept haplotype in all of the isotypes from the Hawaiian Invaded population, particularly on chromosomes I, V, and X, but less so on chromosome IV (**Figure 7B, Supplemental Figure 9**). By contrast, greater than 50% of chromosome IV contained the swept haplotype in all of the isotypes from the global C population (**Supplemental Figure 9)**. Taken together, our data showed that the Hawaiian isotypes from the Volcano, Hawaiian Divergent, and Hawaiian Low populations have avoided the selective sweeps that are pervasive across most regions of the globe, and individuals within the Hawaiian Invaded subpopulation have likely been outcrossed with these swept haplotypes.

## Discussion

We sought to deeply sample the natural genetic variation within the *C. elegans* species to better understand the evolutionary history and driving forces of genome evolution in this powerful model system. Because the Hawaiian Islands have been shown to harbor highly divergent strains relative to most regions of the world, we choose to sample extensively on these islands. We developed a streamlined collection procedure that facilitated our collection of over 2,000 samples across five Hawaiian islands. From these collections, we isolated over 2,500 nematodes and used molecular data to partition 427 of these isolates into 13 distinct taxa, mostly from the *Rhabditidae* family. In total, we identified and cryogenically preserved 95 new *C. elegans* isolates that represent 26 genetically distinct isotypes. These isotypes represent the largest single *C. elegans* collection effort on any island system and contain 27% more SNVs than all 233 non-Hawaiian isotypes combined. Our findings confirm high diversity in Hawaii matching previous studies (Andersen et al., 2012; Wicks et al., 2001). Furthermore, we document the first evidence of outcrossing between Hawaiian and global populations.

### The origins of *C. elegans*

The higher genetic diversity in the Hawaii population might indicate that it represents an ancient population, similar to African populations in humans (Nielsen et al., 2017; Ramachandran et al., 2005). The possibility that the *C. elegans* species might have originated from the Hawaiian Islands, or migrated there from adjacent landmasses shortly after speciation, requires that the Hawaiian Islands predate the split between *C. elegans* and its closest known relative *C. inopinata*, which is estimated to be 10.5 million years (Kanzaki et al., 2018). The extant Hawaii Islands we sampled range in age from the still-forming Big Island to the 5.1 million year old Kauai, but the now submerged Emperor Seamounts represent approximately 70 million years of stable land masses over the Pacific Hotspot (Neall and Trewick, 2008). Therefore, older land masses might have donated colonists to younger islands maintaining the Hawaiian *C. elegans* populations over millions of years and allowing the accumulation of genetic diversity. The higher genetic diversity in the Hawaiian Islands may also be driven by population demography on the Islands. It is possible that Hawaii harbors larger, more temporally stable effective population sizes than other regions of the world that have been sampled. Under a neutral model, populations with a larger effective population size are expected to have a greater number of neutral polymorphisms (Kimura, 1991). These larger, more stable effective population sizes are plausible in Hawaii given the abundant supply of available habitat, *e.g.* rotting fruits and vegetable matter, and stable temperatures throughout the year. The Hawaiian climate is particularly less variable than many temperate regions where *C. elegans* populations must overwinter and are known to exhibit seasonal population expansions and contractions (Frézal and Félix, 2015), and tend to be dominated by highly related genotypes from year to year (Richaud et al., 2018). Ultimately, the pattern of genetic variation in Hawaiian populations is likely influenced by a combination of demographic history (*e.g.*, changes in population size, short-and long-range migration events, and admixture) as well as evolutionary processes such as natural selection, recombination, and mutation. To further untangle the evolutionary history of this species, additional samples from natural areas around the globe and in particular the Pacific Rim will be required.

### Out of Hawaii or invasion of Hawaii?

Our data support outcrossing between the Hawaiian Invaded population and the less-diverse global C population. Moreover, most strains from the Hawaiian Invaded and non-Hawaiian populations share portions of the globally swept haplotype. Within the Hawaiian Invaded population, isotypes share smaller portions of the swept haplotype relative to isotypes from the non-Hawaiian populations. It remains unclear whether sharing of globally swept haplotypes can be explained by emigration of nematodes from Hawaii (out of Hawaii) or immigration of nematodes to Hawaii (invasion of Hawaii). In either case, the Hawaiian Islands are geographically isolated, which should theoretically restrict gene flow to and from the Islands. However, Hawaii’s position as a global trade-hub makes gene flow with the rest of the world more likely (Frankham, 1997). Although we do not have direct evidence to discriminate between these possibilities, the ‘Out of Hawaii’ hypothesis might have occurred through long-range dispersal of genotypes similar to those found in the Hawaiian Invaded population, which then underwent selection over multiple generations to resemble the more swept genotypes found across the globe. Migration out of Hawaii could have been aided by the transition of the Hawaiian economy towards large-scale production and export of sugarcane and tropical fruits, which began in the late nineteenth century (Bartholomew et al., 2012). If correct, then this situation is similar to what is thought to have occurred within *Drosophila melanogaster* where the fruit trade might have facilitated recent migrations from native regions to oceanic islands (David and Capy, 1988; Hales et al., 2015). Alternatively, the pattern of haplotype sharing could be explained by an ‘invasion of Hawaii’ scenario, wherein swept haplotypes have invaded Hawaii. This scenario could threaten the genetic diversity of the Hawaiian populations if the invading alleles confer strong fitness advantages as is expected for swept haplotypes (Andersen et al., 2012). However, if an invasion of Hawaii is currently underway, we have little evidence to support the selection of the globally swept haplotypes in Hawaii. First, the Hawaiian Invaded population only contains small fractions of the swept haplotypes on chromosomes I, V, X, and even smaller fractions on chromosome IV. Second, it would take a considerable number of generations to create the small factions of the swept haplotypes that we observe in the Hawaiian Invaded population because of the low outcrossing rates and high incidence of outbreeding depression in *C. elegans* (Dolgin et al., 2007).

### The ancestral niche of *C. elegans* might be similar to the Hawaiian niche

We used a publicly available weather data from the National Oceanic and Atmospheric Administration and the National Climatic Data Center to measure the variation in seasonal temperatures for locations close to the sites were isotypes were collected (Evans et al., 2017). We found that the Hawaiian populations experienced less seasonal variability in temperature than any of the non-Hawaiian populations (**Supplemental Figure 10**). These findings raise the possibility that the ancestral niche of *C. elegans* might be similar to the thermally stable Hawaiian habitats where genetic diversity is highest. However, factors other than seasonal temperature variation might also characterize the ancestral niche of *C. elegans*. The Hawaiian Divergent population was enriched at higher elevation, which has been less impacted by human activities in Hawaii since the time of Polynesian colonization (Alison Kay, 1994). By contrast, the Hawaiian Invaded population is found at lower elevations. Although it remains unclear what factors restrict gene flow between the non-admixed and Hawaiian Invaded populations, it is possible that selective pressures associated with human impact contribute to their isolation. This possibility would be consistent with the hypothesis that the global sweeps, present in the Hawaiian Invaded population, originated through positive selection acting on loci that confer fitness advantages in human-associated habitats (Andersen et al., 2012). Taken together, we suspect that the ancestral niche of *C. elegans* is likely to be similar to the thermally stable, high elevation Hawaiian habitats where human impacts are less prevalent.

### Unravelling the evolutionary history of *C. elegans*

More accurate models of *C. elegans* niche preferences will facilitate our ability to unravel the evolutionary history of this species by directing researchers to areas most likely to harbor *C. elegans* populations. In order to build more accurate niche models, future sampling efforts should include unbiased sampling across environmental gradients in multiple locations over time because data on niche parameters where *C. elegans* is not found is as important as data where *C. elegans* is found. Additionally, we must identify and quantify important biotic niche factors, including associated bacteria, fungi, and invertebrates. These types of data will help facilitate the identification of genes and molecular processes that are under selection in different subpopulations across the species range. *C. elegans* offers a tractable and powerful animal model system to connect environmental parameters to functional genomic variation. These data will deepen our understanding of the evolutionary history of *C. elegans* by revealing how selection and demographic forces have shaped the genome of this important model system.

## Methods

### Strains

Nematodes were reared at 20°C using OP50 bacteria grown on modified nematode growth medium (NGMA), containing 1% agar and 0.7% agarose to prevent animals from burrowing (Andersen et al., 2014). In total, 169 *C. briggsae*, 100 *C. elegans*, 21 *C. tropicalis*, 15 *C. oiwi*, and four *C. kamaaina* wild isolates were collected. Of these strains, 95 *C. elegans*, 19 *C. tropicalis*, and 12 *C. oiwi* wild isolates were cryopreserved and are available upon request along with the other *C. elegans* strains included in our analysis (**Supplemental File 2)**. The type specimen for *C. oiwi* (ECA1100) is also deposited at the *Caenorhabditis* Genetics Center (**Supplemental File 1**).

### Sampling strategy

We sampled nematodes at 2,263 sites across five Hawaiian Islands during August 2017. Before travelling to Hawaii, general sampling locations were selected based on accessibility via hiking trails and by proximity to where *C. elegans* had been collected previously (Andersen et al., 2012; Cook et al., 2016; Hahnel et al., 2018; Hodgkin and Doniach, 1997). Sampling hikes with large elevation changes were prioritized to ensure that we sampled across a broad range of environmental parameters. On these hikes, we opportunistically sampled substrates known to harbor *C. elegans*, including fruits, nuts, flowers, stems, leaf litter, compost, soil, wood, and live arthropods and molluscs (Ferrari et al., 2017; Frézal and Félix, 2015; Schulenburg and Félix, 2017). In 20 locations, we performed extensive local sampling in an approximately 30 square meter area that we refer to as a ‘gridsect’. The gridsect comprised a center sampling point with additional sampling sites at one, two, and three meters away from the center in six directions with each direction 60° apart from each other (**Supplemental Figure 1**).

### Field sampling and environmental data collection

To characterize the *Caenorhabditis* abiotic niche, we collected and organized data for several environmental parameters at each sampling site using a customizable geographic data-collection application called Fulcrum®. We named our customized Fulcrum® application ‘Nematode field sampling’ and used the following workflow to enter the environmental data into the application while in the field. First, we used a mobile device camera to scan a unique collection barcode from a pre-labelled plastic collection bag. This barcode is referred to as a collection label or ‘C-label’ in the application and is used to associate a particular sample with its environmental and nematode isolation data. Next, we entered the substrate type, landscape, and sky view data into the application using drop down menus and photographed the sample in place using a mobile device camera. The GPS coordinates for the sample are automatically recorded in the photo metadata. We then measured the surface temperature of the sample using an infrared thermometer Lasergrip 1080 (Etekcity, Anaheim, CA), its moisture content using a handheld pin-type wood moisture meter MD912 (Dr. Meter, Los Angeles, CA), and the ambient temperature and humidity near the sample using a combined thermometer and hygrometer device GM1362 (GoerTek, Weifang, China). These measurements were entered into the appropriate fields in the application (**Supplemental Table 3**). Finally, we transferred the sample into a collection bag and stored it in a cool location before we attempted to isolate nematodes. Seventy samples in our raw data had missing GPS coordinates or GPS coordinates that were distant from actual sampling locations after visual inspection using satellite imagery. The positions for these samples were corrected using the average position of the two samples collected before and after the errant data point or by manually assigning estimated positions.

### Nematode isolation

Following each collection, the substrate sample was transferred from the barcoded collection bag to an identically barcoded 10 cm NGMA plate seeded with OP50 bacteria. For 1,989 of the 2,263 samples collected, we isolated nematodes that crawled off the substrates onto the collection plates approximately 47 hours after the samples were collected from the field (mean = 46.9 h, std. dev. = 19.5 h). The remaining 274 samples were shipped overnight from Hawaii to Northwestern University in collection bags, and the nematodes were isolated approximately 172 hours after sample collection (mean = 172.5 h, std. dev. = 17.9 h). For each collection plate, up to seven gravid nematodes were isolated by transferring them individually to pre-labeled 3.5 cm NGMA isolation plates seeded with OP50 bacteria. We refer to these isolation plates as ‘S-plates’ in the Fulcrum® application we called ‘Nematode isolation’ (**Supplemental Table 4)**. At the time of isolation, we recorded the approximate number of nematodes on the collection plate and whether males or dauers were present. Importantly, male and dauer observations from samples shipped from Hawaii were not recorded to avoid bias caused by the long handling time of these samples. We merged the collection, isolation, and environmental data together into a single data file with the ‘process_fulcrum_data.R’ script that can be found in the scripts folder of the GitHub repo (https://github.com/AndersenLab/Hawaii_Manuscript) (**Supplemental Data 4**).

### Nematode identification

The isolated nematodes were stored at 20°C for approximately 14 days (mean = 14.3 d, std. Dev. = 4.9 d) but were not passaged during this time to avoid multiple generations of proliferation. For initial genotyping, five to ten nematodes were lysed in 8 µl of lysis solution (100 mM KCl, 20 mM Tris pH 8.2, 5 mM MgCl_2_, 0.9% IGEPAL, 0.9% Tween 20, 0.02% gelatin with proteinase K added to a final concentration of 0.4 mg/ml) then frozen at −80°C for up to 12 hours. The lysed material was thawed on ice, and 1 µl was loaded directly into 40 µl reactions with primers spanning a portion of the ITS2 region (Internal Transcribed Spacer) between the 5.8S and 28S rDNA genes with forward primer oECA305 (GCTGCGTTATTTACCACGAATTGCARAC) and reverse primer oECA202 (GCGGTATTTGCTACTACCAYYAMGATCTGC) (Kiontke et al., 2011). The PCR used the following conditions: three minutes denaturation step at 95°C; then 34 cycles of 95°C for 15 seconds, 60°C for 15 seconds, and 72°C for two minutes; followed by a five-minute elongation step at 72°C. The presence of ITS2 PCR products was visualized on a 2% agarose gel in 1X TAE buffer. Isolates that did not yield an ITS2 PCR product were labelled as ‘PCR-negative’, and those reactions that yielded the expected 2 kb ITS2 PCR product were labelled as ‘PCR-positive’. We then used Sanger sequencing to sequence the ITS2 PCR products with forward primer oECA305. We classified *Caenorhabditis* species by comparing the ITS2 sequences to the National Center for Biotechnology Information (NCBI) database using the BLAST algorithm. Isolates with sequences that aligned best to genera other than *Caenorhabditis* were only classified to the genus level. For every isolate where the BLAST results either aligned to *C. elegans*, had an unexpectedly high number of mismatches in the center of the read, or did not match any known sequences because of poor sequence quality, we performed another independent lysis and PCR using high-quality Taq polymerase (cat# RR001C, TaKaRa) to confirm our original results. For this confirmation, we used the forward primer oECA305 and the reverse primer oECA306 (CACTTTCAAGCAACCCGAC) to sequence the confirmation ITS2 amplicon in both directions. The sequence chromatograms were then quality trimmed by eye with Unipro UGENE software (version 1.27.0) and compared to known nematode species in the NCBI sequence database using the BLAST algorithm. We used the consensus alignment of the forward and reverse reads to confirm our original results. For *C. elegans*, five of the 100 strains perished before we could confirm their identity. We also confirmed that several strains that best aligned to *C. kamaaina* shared a large number of mismatches in the center of the ITS2 amplicon, suggesting they belonged to a new species. For these strains, we performed reciprocal mating tests with *C. kamaaina* to infer the new species by the biological species concept (Félix et al., 2014). None of these crosses produced viable progeny, suggesting that these isolates represent a new *Caenorhabditis* species (**Supplemental File 1**).

### Illumina library construction and whole-genome sequencing

To extract DNA, we transferred nematodes from two 10 cm NGMA plates spotted with OP50 *E. coli* into a 15 ml conical tube by washing with 10 mL of M9. We then used gravity to settle animals on the bottom of the conical tube, removed the supernatant, and added 10 mL of fresh M9. We repeated this wash method three times over the course of one hour to serially dilute the *E. coli* in the M9 and allow the animals time to purge ingested *E. coli*. Genomic DNA was isolated from 100-300 µl nematode pellets using the Blood and Tissue DNA isolation kit cat# 69506 (QIAGEN, Valencia, CA) following established protocols (Cook et al., 2016). The DNA concentration was determined for each sample with the Qubit dsDNA Broad Range Assay Kit cat# Q32850 (Invitrogen, Carlsbad, CA). The DNA samples were then submitted to the Duke Center for Genomic and Computational Biology per their requirements. The Illumina library construction and sequencing were performed at Duke University using KAPA Hyper Prep kits (Kapa Biosystems, Wilmington, MA) and the Illumina NovaSeq 6000 platform (paired-end 150 bp reads). The raw sequencing reads for strains used in this project are available from the NCBI Sequence Read Archive (Project PRJNA549503).

### Variant calling

To ensure reproducible data analysis, all genomic analyses were performed using pipelines generated in the Nextflow workflow management system framework (Di Tommaso et al., 2017). Each Nextflow pipeline used in this study is briefly described below (**Supplemental Table 5**). All pipelines follow the “*pipeline name-nf*” naming convention and full descriptions can be found on the Andersen lab dry-guide website: (http://andersenlab.org/dry-guide/pipeline-overview/).

Raw sequencing reads were trimmed using *trimmomatic-nf*, which uses trimmomatic (v0.36) (Bolger et al., 2014) to remove low-quality bases and adapter sequences. Following trimming, we used the *concordance-nf* pipeline to characterize *C. elegans* strains isolated in this study and previously described strains (Cook et al., 2017, 2016; Hahnel et al., 2018). The *concordance-nf* pipeline calls SNVs using the BCFtools (v.1.9) (Danecek et al., 2014) variant calling software. The variants are filtered by: Depth (FORMAT/DP) ≥ 3; Mapping Quality (INFO/MQ) > 40; Variant quality (QUAL) > 30; (Allelic Depth (FORMAT/AD) / Num of high quality bases (FORMAT/DP)) ratio > 0.5. We determined the pairwise similarity of all strains by calculating the fraction of shared SNVs. Finally, we classified two or more strains as the same isotype if they shared >99.9% SNVs. If a strain did not meet this criterion, we considered it as a unique isotype. Newly assigned isotypes were added to CeNDR (Cook et al., 2017).

After isotypes are assigned, we used *alignment-nf* with BWA (v0.7.17-r1188) (Li, 2013; Li and Durbin, 2009) to align trimmed sequence data for distinct isotypes to the N2 reference genome (WS245) (Lee et al., 2018). Next, we called SNVs using *wi-nf*, which uses the BCFtools (v.1.9) (Danecek et al., 2014). The *wi-nf* pipeline generates two population-wide VCFs that we refer to as the soft-filtered and hard-filtered VCFs (**Supplemental Table 2**). After variant calling, a soft-filtered VCF was generated for each sample by appending the following soft-filters to variant sites: Depth (FORMAT/DP) > 10; Mapping Quality (INFO/MQ) > 40; Variant quality (QUAL) > 10; (Allelic Depth (FORMAT/AD) / Number of high quality bases (FORMAT/DP)) ratio > 0.5. These soft-filters were appended to the FT field of the VCF using *VCF-kit* (Cook and Andersen, 2017). Next, sample VCFs were merged using the merge utility of BCFtools. Once the population VCF was generated, variant sites with greater than 90% missing genotypes (high_missing) or greater than 10% heterozygosity (high_heterozygosity) were flagged. We refer to this VCF as the soft-filtered VCF. To construct the hard-filtered VCF, we removed all variants that did not pass the filters described above. Both the soft-and hard-filtered isotype-level VCFs are available to download on the CeNDR website (version 20180527) (Cook et al., 2017).

We further pruned the hard-filtered VCF to contain sites with no missing genotype calls and removed sites in high linkage disequilibrium (LD) using PLINK (v1.9) (Chang et al., 2015; Purcell et al., 2007) with the*--indep-pairwise 50 1 0.95* command. The predicted variant effects were appended to the VCF using SnpEff (v 4.3) (Cingolani et al., 2012). We further annotated this VCF with exons, G-quartets, transcription factor binding sites, histone binding sites, miRNA binding sites, splice sites, ancestral alleles (XZ1516 set as ancestor), the genetic map position, and repetitive elements using vcfanno (v 0.2.8) (Pedersen et al., 2016). All annotations were obtained from WS266. We removed regions that were annotated as repetitive. We named this VCF the ‘PopGen VCF’ (**Supplemental Data 4; Supplemental Table 2**).

### Phylogenetic analyses

We characterized the relatedness of the *C. elegans* population using RAxML-ng with the GTR DNA substitution model and maximum likelihood estimation to find the parameter values that maximize the phylogenetic likelihood function, and thus provide the best explanation for the observed data (Kozlov et al., 2019). We used the vcf2phylip.py script (Ortiz, n.d.) to convert the ‘PopGen VCF’ (**Supplemental Data 4**) to the PHYLIP format (Felsenstein, 1993) required to run RAxML-ng. To construct the tree that included 276 strains, we used the GTR evolutionary model available in RAxML-ng (Lanave et al., 1984; Tavaré, 1986). Trees were visualized using the ggtree (v1.10.5) R package (Yu et al., 2017). To construct the neighbor-net phylogeny, we used SplitsTree4 (Huson and Bryant, 2006).

### Population genetic statistics

Genome-wide pi, Hudson’s *F*_*ST*_, and Tajima’s D were calculated using the PopGenome package in R (Pfeifer et al., 2014). All statistics were calculated along sliding windows with a 10 kb window size and a 1 kb step size.

### Admixture analysis

We performed admixture analysis using ADMIXTURE (v1.3.0) (Alexander et al., 2009). Prior to running ADMIXTURE, we LD-pruned the ‘PopGen VCF’ (**Supplemental Data 4**) using PLINK (v1.9) (Chang et al., 2015; Purcell et al., 2007) with the command *--indep-pairwise 50 10 0.8*. We also removed variants only present in one isotype. We ran ADMIXTURE ten independent times for K sizes ranging from 2 to 20 for all 276 isotypes. Visualization of admixture results was performed using the pophelper (v2.2.5) R package (Francis, 2017). We chose K=11 for future analyses because the cross-validation (CV) error approached minimization at this K (**Supplemental Figure 5**). Furthermore, K=11 subset the Hawaiian isotypes into four distinct populations, which exactly matched the subsets obtained from running ADMIXTURE on just the 43 Hawaiian isotypes at K=4 (K=4 minimized CV for ADMIXTURE with Hawaiian isotypes only, (**Supplemental Figure 11**). We performed TreeMix analysis on K=11 for zero to five migration events (Pickrell and Pritchard, 2012).

### Haplotype analysis

We determined identity-by-descent (IBD) of strains using IBDSeq (Browning and Browning, 2013) run on the ‘PopGen VCF’ (**Supplemental Data 4**) with the following parameters: *minalleles=0.01, ibdtrim=0, r2max=0.8*. IBD segments were then used to infer haplotype structure among isotypes as described previously (Andersen et al., 2012). After haplotypes were identified, we defined the most common haplotype found on chromosomes I, IV, V, and X as the swept haplotype. We then retained the swept haplotypes within isotypes that passed the following per chromosome filters: total length > 1 Mb; total length / maximum population-wide swept haplotype length > 0.03. We classified chromosomes within isotypes as swept if the sum of the retained swept haplotypes for a chromosome was > 3% of the maximum population wide swept haplotype length for that chromosome.

### Environmental parameter analysis

We calculated the pairwise distances among all *C. elegans*-positive collections on Hawaii and detected five distinct geographic clusters, each of which contain collections that are within 20 meters of one another. The largest of these clusters comprised 18 collections in the Kalopa State Recreation Area on the Big Island of Hawaii. This cluster contained 11 collections from gridsect-3 and seven additional collections within 20 meters from the edge of the gridsect. The other four geographic clusters contain four or fewer collections each. We used the average values of environmental parameters from geographically clustered collections to avoid biasing our results by local oversampling. We applied this strategy to the comparison of environmental parameters between the Hawaiian admixture populations and used the Kuskal-Wallis test to detect differences (α = 0.05).

## Acknowledgements

We thank the members of the Andersen lab for editing the manuscript for flow and content and for making reagents used in the experiments presented. We are grateful to landowners who gave us permission to collect nematodes on their property. We also thank individuals who have helped us collect additional strains. We would also like to thank the Hawaii Department of Land and Natural Resources as well as the Natural Area Reserves System for permitting, support for these studies, and general advice about the Hawaiian Islands. Additionally, Dr. Sam Gon from The Nature Conservancy Hawai’i Program helped with the naming of *Caenorhabditis oiwi*. This research was supported by start-up funds from Weinberg College of Arts and Sciences and the Molecular Biosciences department. KK is supported by NSF DEB 0922012 to D. H. A. Fitch.

