## Supplemental Figures and Tables for "Deep sampling of Hawaiian *Caenorhabditis elegans* reveals high genetic diversity and admixture with global populations"

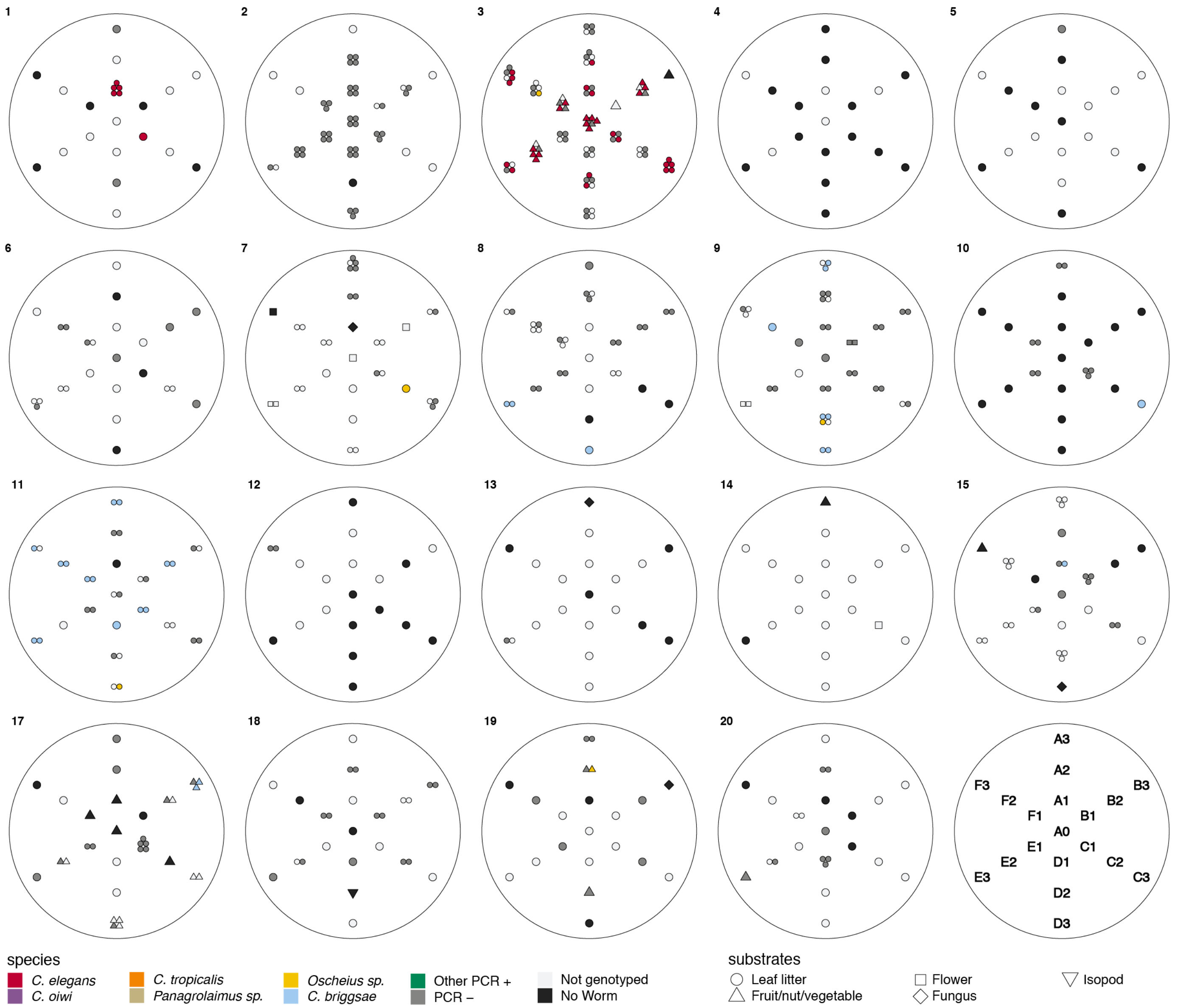


**Supplemental Figure 1 - Local scale sampling with gridsects.** Nineteen of twenty gridsects are shown. A key defining the sample positions for each gridsect is shown on the lower right. Samples were collected in six directions (A-F), each 60º apart. The center is denoted as A0, and samples were collected in each direction at one, two, and three meters from the center. Gridsect 16 is omitted because it was incomplete. The colors and shapes plotted at gridsect positions show the collection category and substrate class collected at that position as defined in the plot legend. We isolated multiple nematodes from some substrates, in these cases multiple shapes are plotted at that position to indicate the various collection categories that were found there.

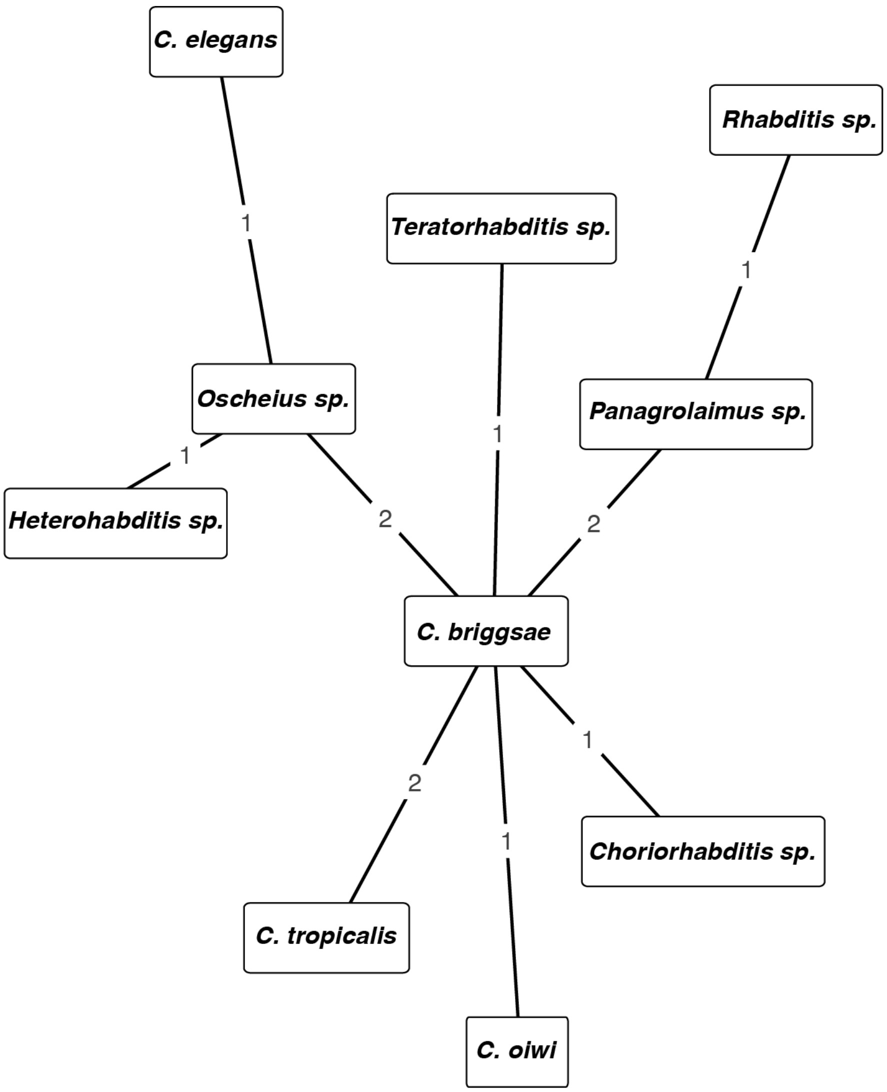


**Supplemental Figure 2 - Network of cohabiting species isolated from samples**. The nodes are labeled with the taxa. The edges are labeled with the number of times the taxa shown on the nodes were isolated from the same sample.

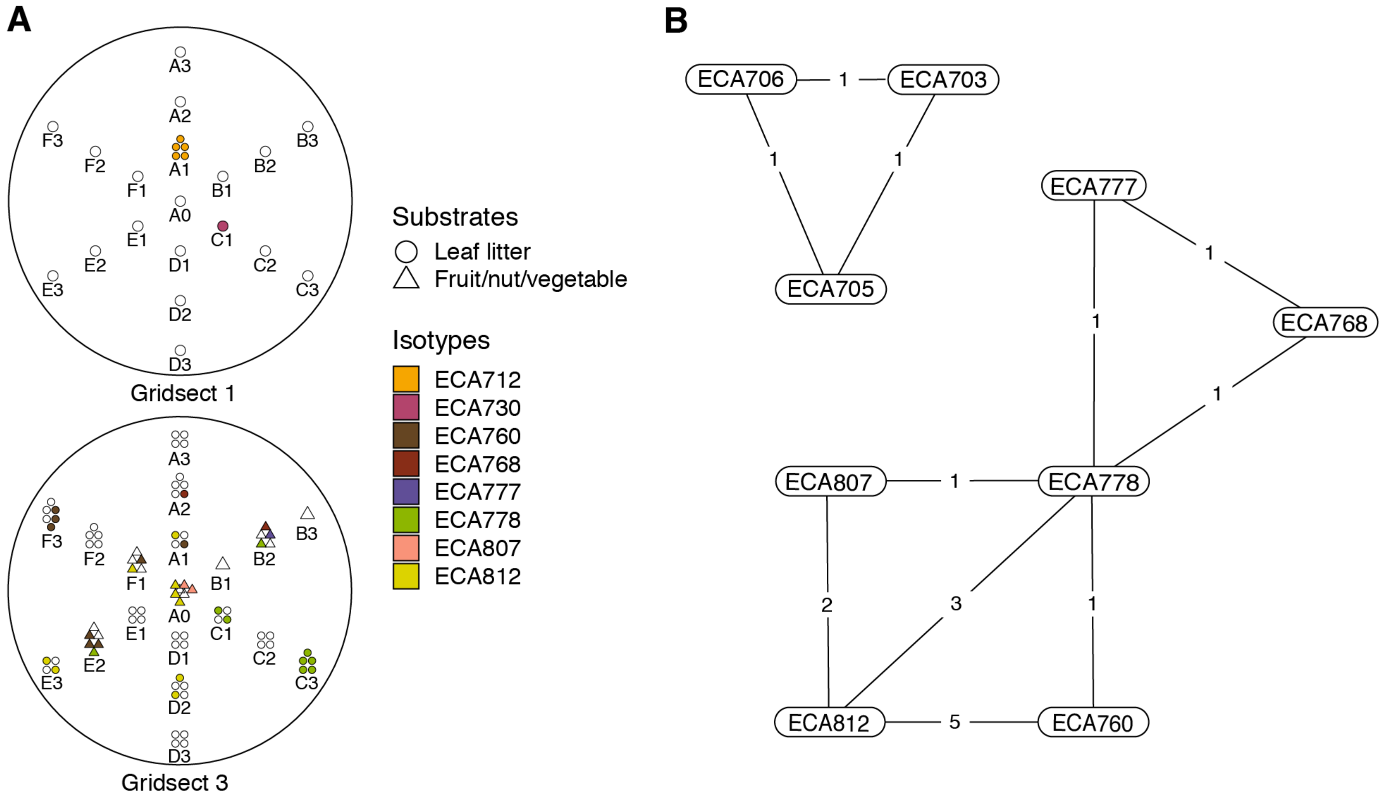


**Supplemental Figure 3 - Local diversity and colocalization of isotypes.** (**A**) Local scale gridsect sampling is shown. Each gridsect contains a total of 19 samples centered on the sample position labeled A0. The remaining sample positions are labeled by one of six transect lines (A-F) followed by the distance (in meters) from position A0. The shapes plotted above the position label show the collection category at that position as defined in the plot legend, and the colors correspond to *C. elegans* isotypes in the legend. Samples that did not contain *C. elegans* strains are colored white. (**B**) A colocalization network is shown for *C. elegans* isotypes from all Hawaiian samples. The numbers inset on the lines connecting two isotypes correspond to the number of unique samples where the two strains were isolated together.

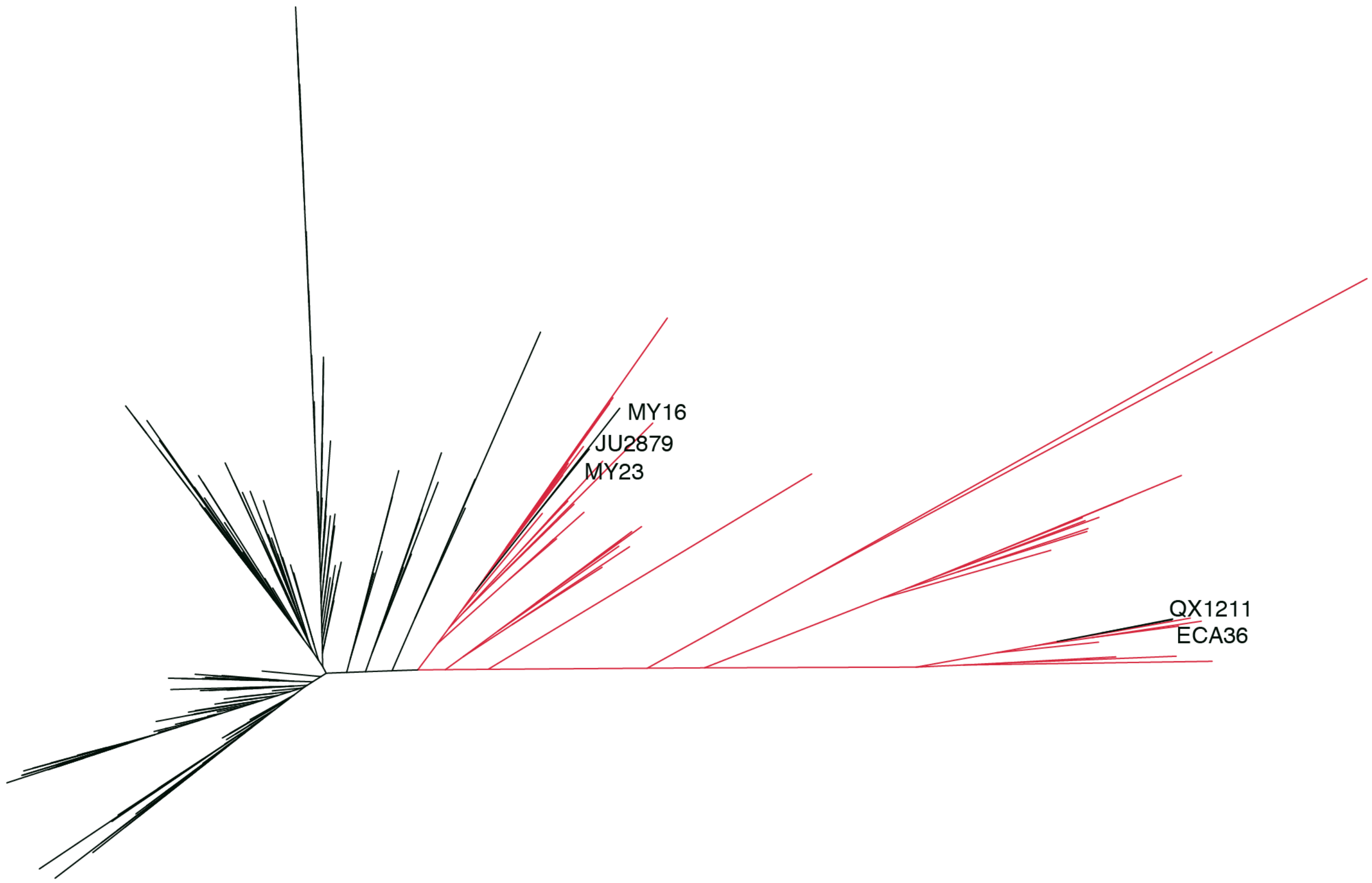


**Supplemental Figure 4 - *Caenorhabditis elegans* unrooted tree for 276 isotypes.** A maximum likelihood tree built using single nucleotide variants found in the 276 isotype *C. elegans* population, including the 26 new Hawaiian isotypes. (Substitution model: GTR+FO). The isotypes labeled in red were isolated from the Hawaiian Islands (n = 43). The five isotypes labeled are non-Hawaiian strains that group with the Hawaiian isotypes.

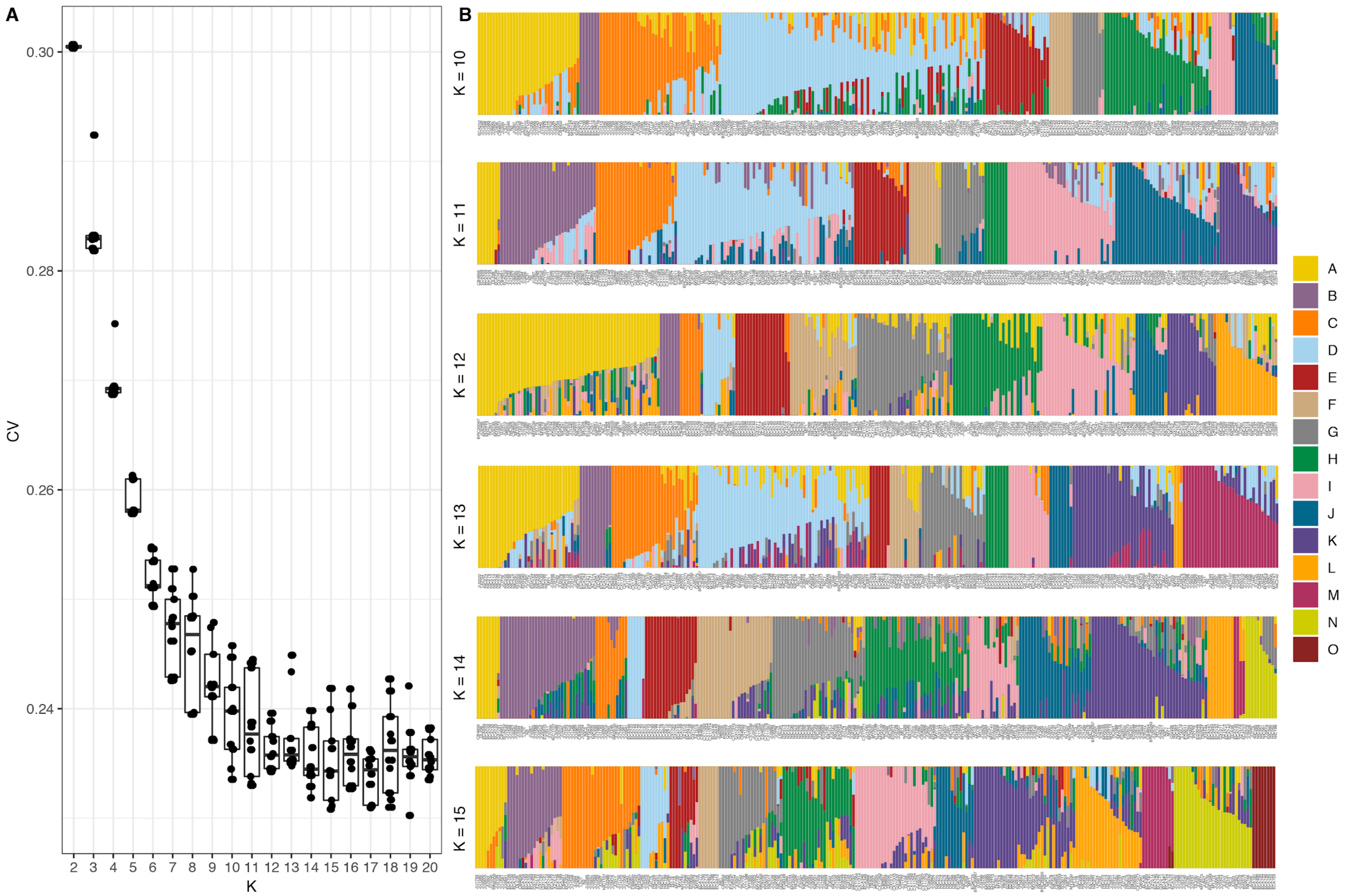


**Supplemental Figure 5 - Summary of ADMIXTURE analysis**. (**A**) Tukey boxplots of ten independent ADMIXTURE runs showing the cross-validation error on the y axis for the ancestral population sizes ranging from 2-20 on the x-axis. (**B**) The inferred ancestral population proportions estimated by ADMIXTURE are shown on the y-axis of the all *C. elegans* strains on the x-axis. The colors correspond to inferred ancestral populations. We have not specified isotype names in the figure because population names are not consistent across Ks.

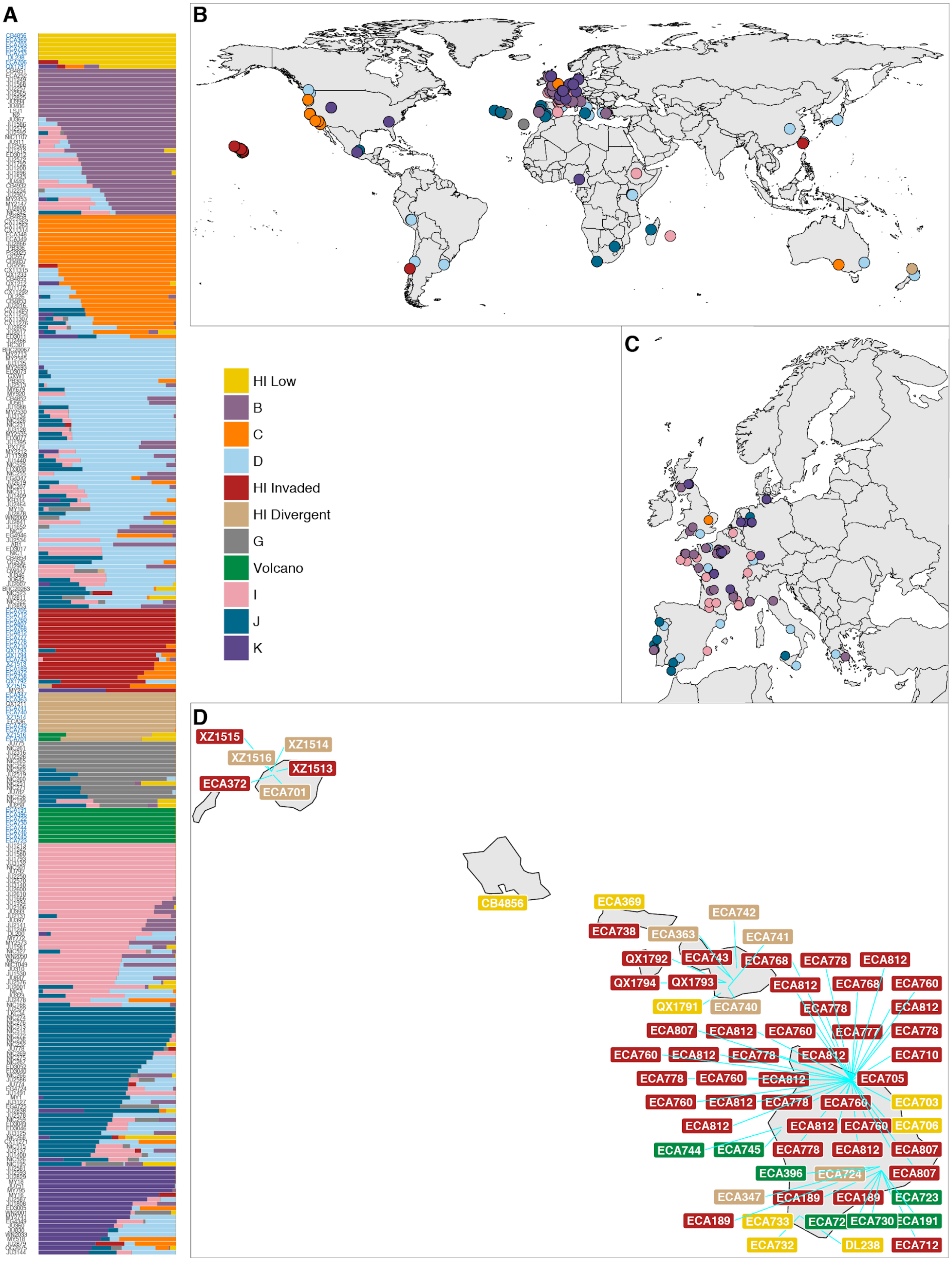
A
B
C
D
HI Low
B
C
D
HI Invaded
HI Divergent
G
Volcano
I
J
K
XZ1515
XZ1514
XZ1516
XZ1513
ECA372
ECA701
CB4856
ECA369
ECA742
ECA738
ECA363
ECA741
QX1792
ECA743
ECA768
ECA778
ECA812
QX1794
QX1793
ECA812
ECA768
ECA760
QX1791
ECA740
ECA778
ECA812
ECA807
ECA812
ECA760
ECA777
ECA778
ECA760
ECA812
ECA778
ECA812
ECA710
ECA778
ECA760
ECA812
ECA705
ECA760
ECA812
ECA778
ECA760
ECA703
ECA812
ECA812
ECA760
ECA706
ECA744
ECA745
ECA778
ECA812
ECA807
ECA396
ECA724
ECA807
ECA347
ECA189
ECA189
ECA723
ECA189
ECA733
ECA72
ECA730
ECA191
ECA732
DL238
ECA712

**Supplemental Figure 6 - Global population structure**. (**A**) The inferred ancestral population proportions estimated by ADMIXTURE (K = 11) with *C. elegans* isotype names on the y-axis. The names colored in blue represent Hawaiian isotypes. (**B**) The global distribution of all 276 isotypes are shown with colors corresponding to the legend. Colors are assigned based on the largest ancestral population fraction for that isotype (*e.g.* an isotype assigned as 60% global C and 40% global D will be colored orange for global C). (**C**) The same data are shown but with more resolution in Europe. (**D**) All distinct collections of Hawaiian isotypes are shown with labels for isotype names colored by the ancestral population assignment. The blue lines point to specific collection locations for those isotypes. The cluster of Hawaiian Admixed isotypes on the Big Island correspond to gridsect-3 and adjacent collections from the Kalōpā state recreation area.

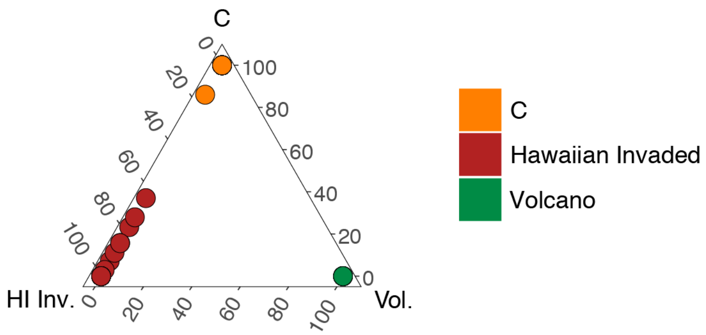


**Supplemental Figure 7 - Admixture between the Hawaiian Invaded and global C populations**. The fraction of admixture among the three populations is shown on a ternary plot. The data points correspond to isotypes and are colored by the highest fractional population assignment.

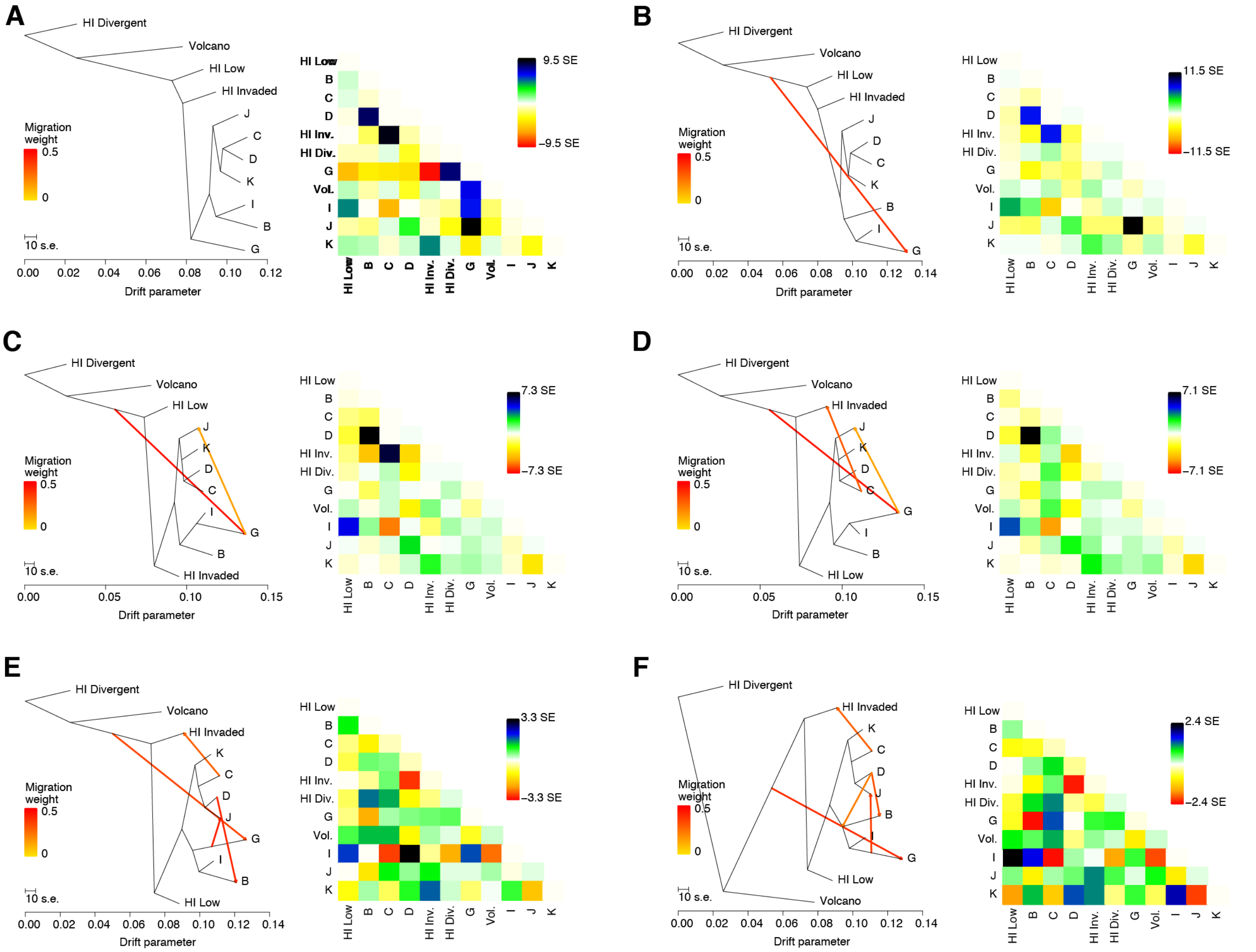


**Supplemental Figure 8 - Evidence of migration between Hawaiian Invaded and global C population.** (**A-D**) The inferred relationship among the ancestral populations (ADMIXTURE, K=11) with zero (**A**), one (**B**), two (**C**), three (**D**), four (**E**), or five (**F**) migration events. The right panels contain heat maps corresponding to the residual fit to the migration models.

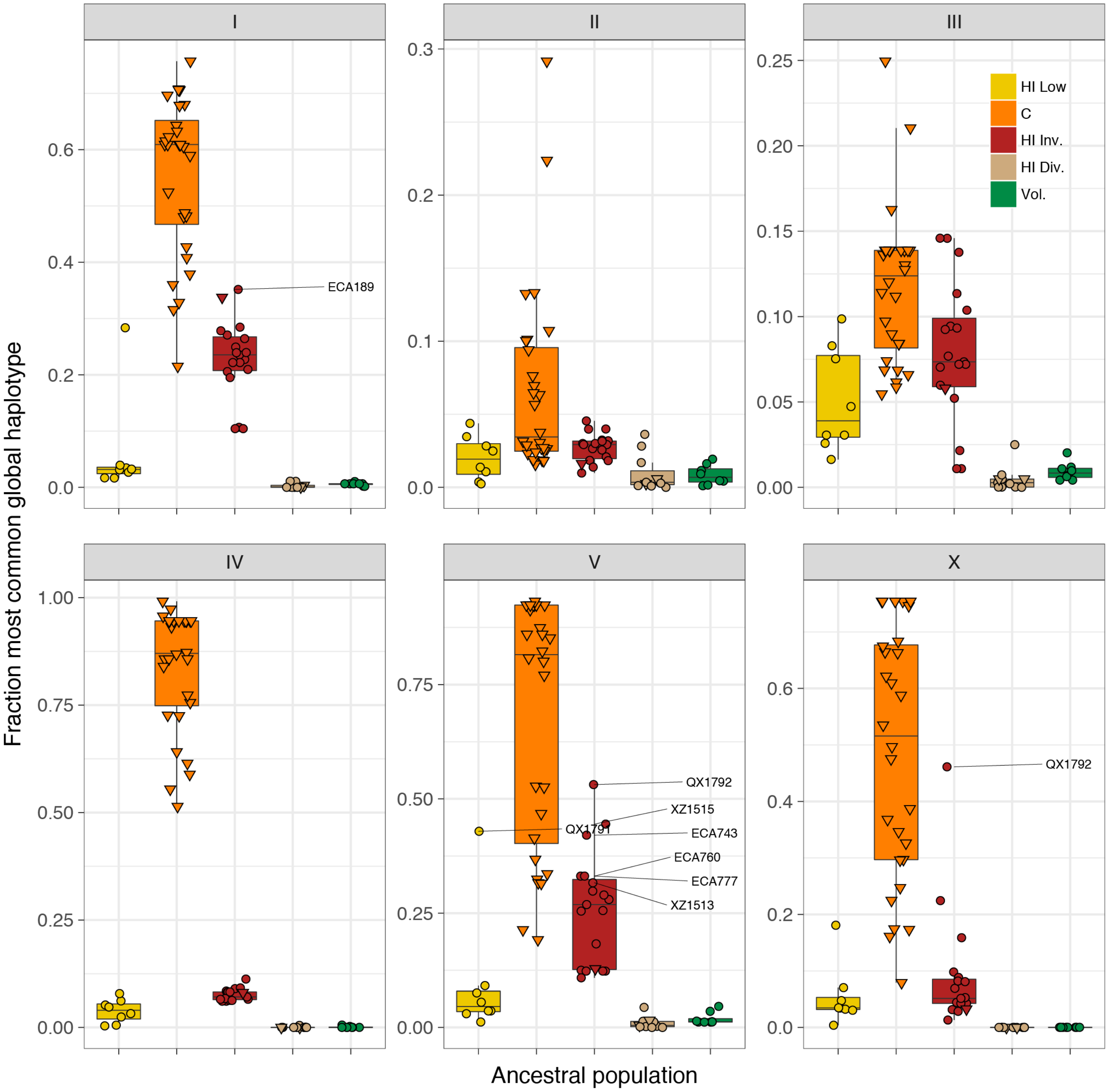


**Supplemental Figure 9 - Most common global haplotype sharing by chromosome and ancestral population.** The fraction of each chromosome that belongs to the most common global haplotype is shown on the y axis. The data points correspond to isotypes and are colored by their assigned ancestral populations. The Hawaiian isotypes are plotted as circles and non-Hawaiian isotypes are plotted as triangles. Hawaiian isotypes with greater than 30% of a chromosome belonging to the most common global haplotype are labelled in the chromosome facet. The most common global haplotype is synonymous with the globally swept haplotype on chromosomes I, IV, V, and the left of X.

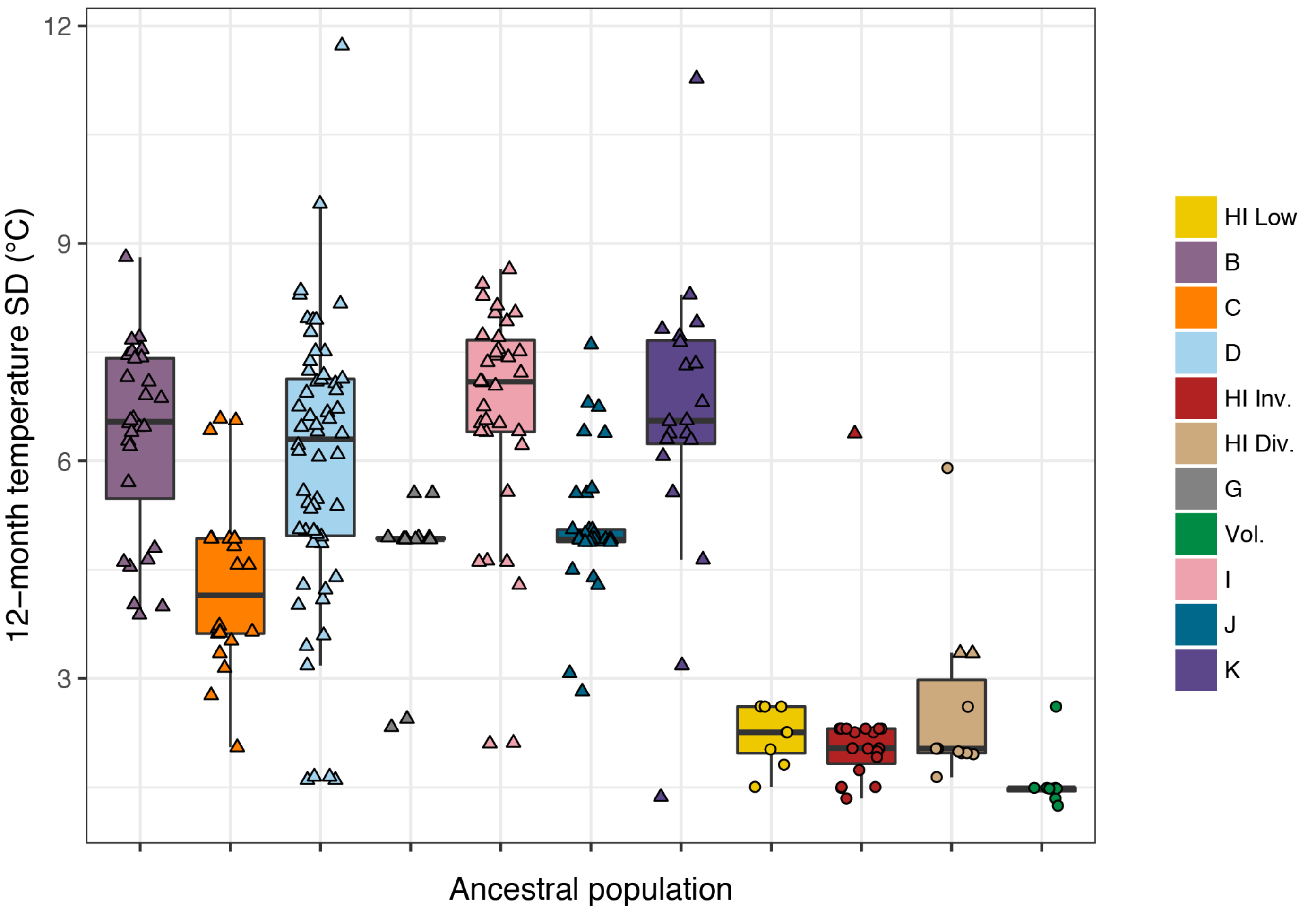


**Supplemental Figure 10 - Seasonal temperature variation by ancestral population.** The standard deviation of daily mean temperatures for a 12-month period centered on the collection date for each isolate is shown. If only the year of collection is known, then the 12-month period is centered on January 1st of that year, and if the year and month are known but not the exact date, then the 12-month period is centered on the first of that month. The data points correspond to isotypes and are colored by their assigned ancestral populations from admixture analysis. The Hawaiian isotypes are plotted as circles and non-Hawaiian isotypes are plotted as triangles.

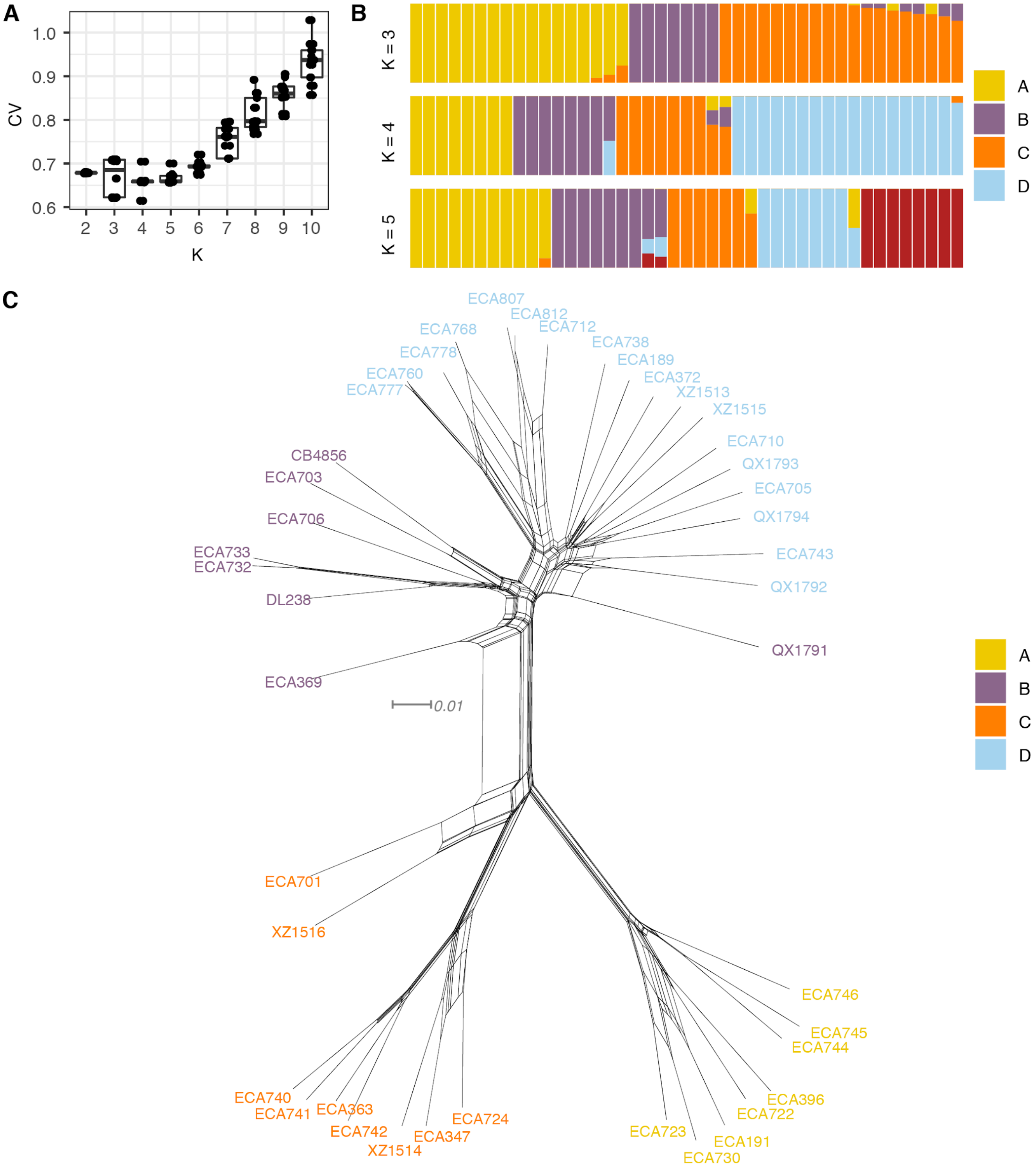


**Supplemental Figure 11 - Summary of ADMIXTURE analysis on Hawaiian *C. elegans* isotypes**. (**A**) Tukey boxplots of ten independent ADMIXTURE runs showing the cross-validation error on the y-axis for the ancestral population sizes ranging from 2-10 on the x-axis. (**B**) The inferred ancestral population proportions estimated by ADMIXTURE are shown on the y-axis of the Hawaiian *C. elegans* isotypes on the x-axis. The colors correspond to inferred ancestral populations. We have not specified isotype names in the figure because population names are not consistent across Ks. (**C**) A neighbor-joining net showing the genetic relatedness of the Hawaiian *C. elegans* population is shown. Colors of labels indicate the ancestral population assignment from ADMIXTURE (K=4).

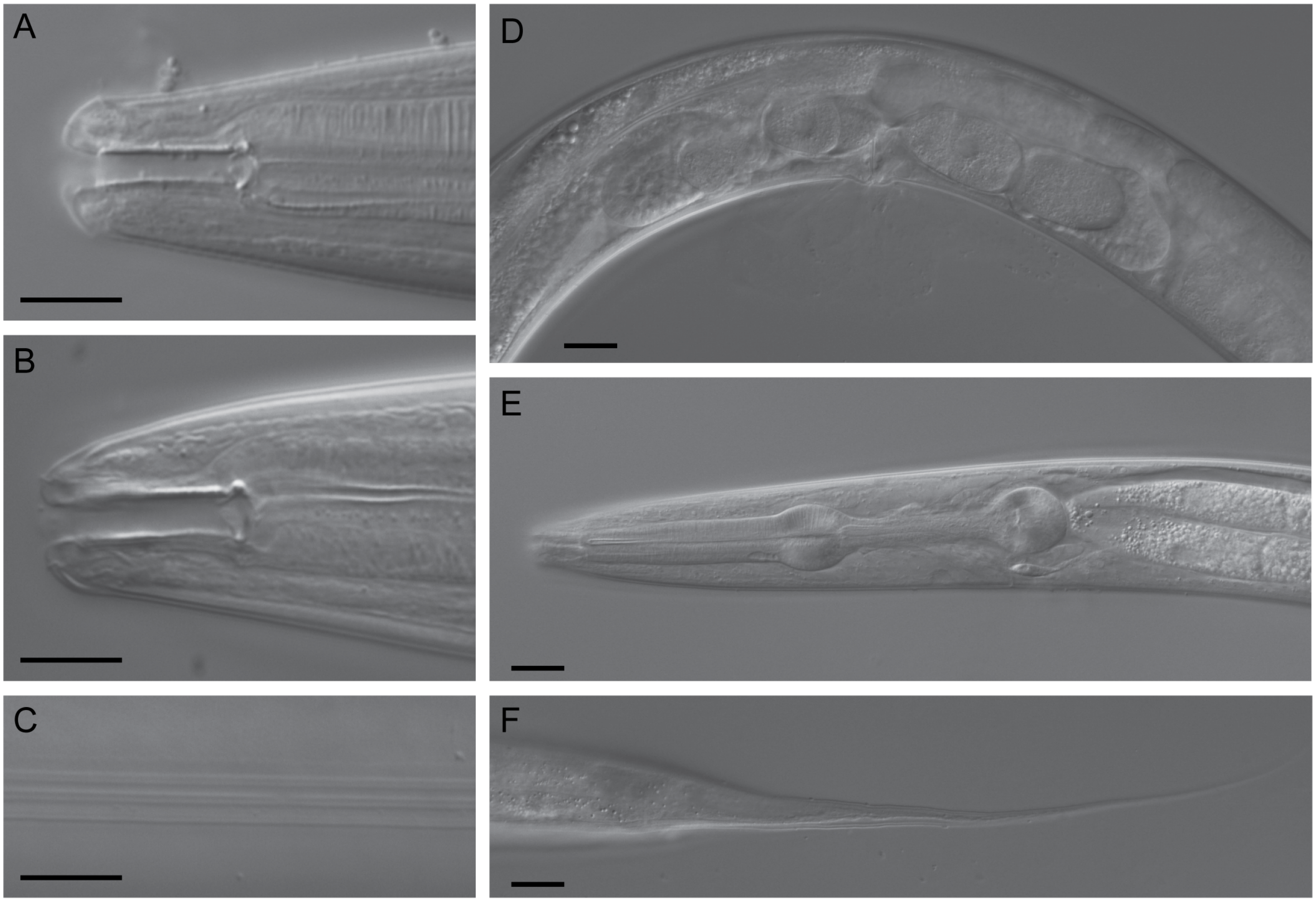


**Supplemental Figure 12 - DIC micrographs of *C. oiwi* sp. n.** (**A**) stoma of male (subventral right, dorsal is up); (**B**) stoma of female subventral right, dorsal is up); (**C**) female lateral field with alae; (**D**) female midbody region showing vulva, one embryo in each uterus, one oocyte in each spermatheca and part of the posterior ovary (left side view); (**E**) pharynx region of female (left side view); and (**F**) female tail (left side view). Scale bars in A-C are 10 μm and 20 μm in D-F.

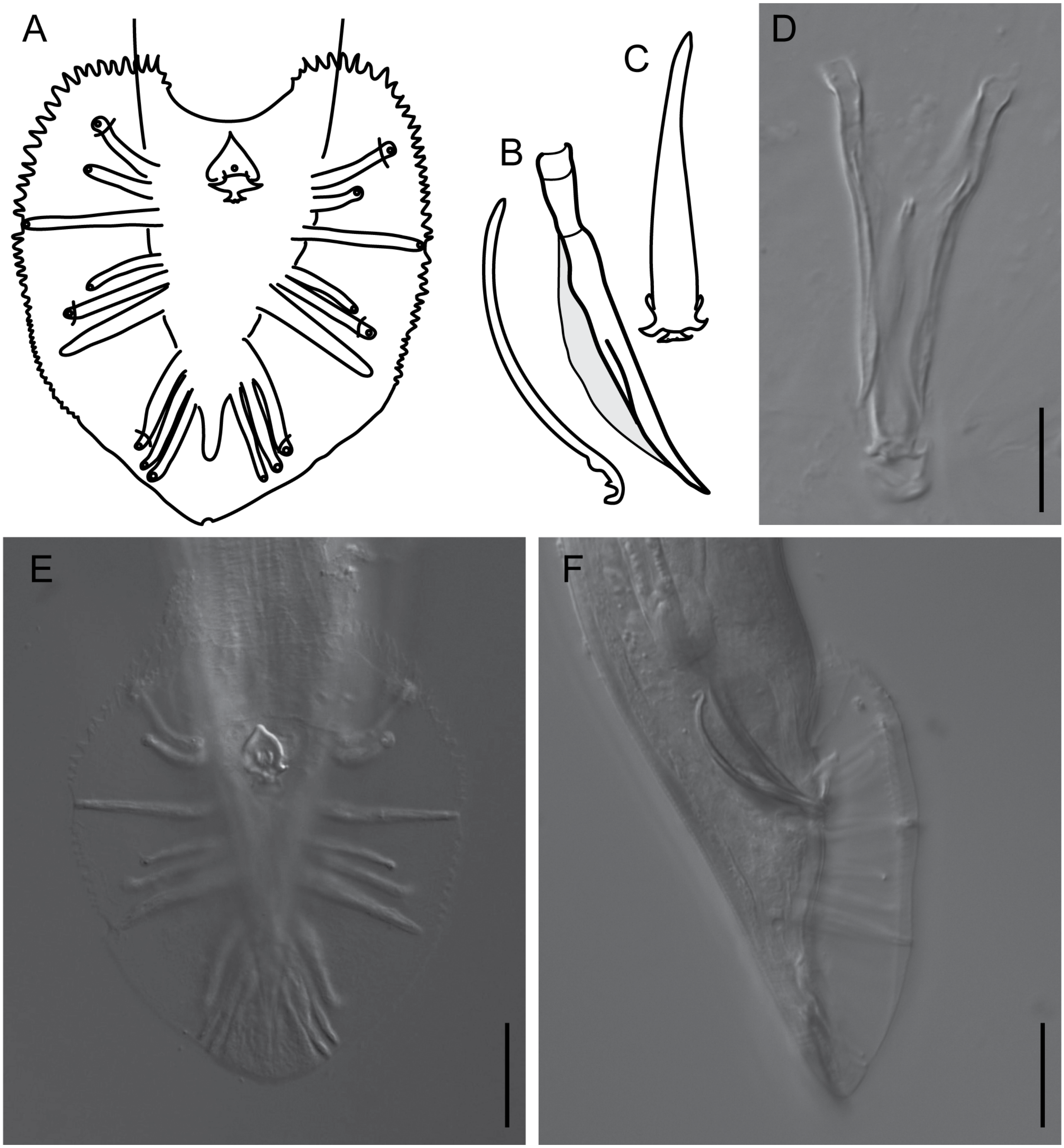


**Supplemental Figure 13 - Features of the male tail of *C. oiwi* sp. n.** (**A**) Drawing of the male tail in ventral view. The rays in position 1, 5 and 7 (from anterior) open to the dorsal side of the fan. (**B**) A drawing of the spicule and guernaculum in right lateral view is shown. (**C**) A drawing of the gubernaculum in ventral view is shown. (**D**) DIC micrograph of the spicules and gubernaculum in ventral view is shown. (**E-F**) DIC micrographs of the male tail in ventral (**E**) and lateral right view (**F**) are shown. Scale bars are 20 µm.

| **Collection Category** | **Big Island** | **Kauai** | **Maui** | **Molokai** | **Oahu** | **Total** |
|---|---|---|---|---|---|---|
| *C. elegans* | 31 | 1 | 5 | 1 | 0 | 38 |
| *C. oiwi* | 1 | 3 | 0 | 0 | 8 | 12 |
| *C. tropicalis* | 5 | 3 | 3 | 0 | 2 | 13 |
| *C. kamaaina* | 0 | 2 | 0 | 0 | 0 | 2 |
| *C. briggsae* | 20 | 49 | 22 | 0 | 4 | 95 |
| *Panagrolaimus* sp. | 7 | 1 | 3 | 1 | 0 | 12 |
| *Oscheius* sp. | 25 | 19 | 7 | 1 | 6 | 58 |
| *Teratorhabditis* sp. | 0 | 0 | 1 | 0 | 0 | 1 |
| *Rhabditis* sp. | 3 | 0 | 1 | 0 | 0 | 4 |
| *Choriorhabditis* sp. | 0 | 1 | 2 | 0 | 0 | 3 |
| *Mesorhabditis* sp. | 1 | 0 | 0 | 0 | 0 | 1 |
| *Chabertia* sp. | 1 | 0 | 0 | 0 | 0 | 1 |
| *Heterorhabditis* sp. | 1 | 0 | 0 | 0 | 0 | 1 |
| PCR - | 272 | 246 | 111 | 22 | 32 | 683 |
| Not genotyped | 343 | 296 | 239 | 42 | 38 | 958 |
| No Worm | 124 | 407 | 129 | 23 | 29 | 712 |
| Total | 834 | 1028 | 523 | 90 | 119 | 2594 |

**Supplemental Table 1 - Collection categories identified on each island.** Multiple collection categories were found for some samples. For this reason, the total number of distinct collections (2,594) exceeds the total number of samples (2,263).

| VCF | Description | Use | Filter Parameters | Availability | Software used |
|---|---|---|---|---|---|
| soft-filtered | Unfiltered variant set with soft-filters appended at variant and sample level | initial variant calling | Depth (DP) > 10;<br>Mapping Quality (MQ) > 40;<br>Variant quality (QUAL) > 10;<br>((AD) / (DP)) ratio > 0.5;<br>high_heterozygosity: >10% het calls ; high_missing > 90% sites missing | CeNDR | BCFtools - variant calling and variant level filters; vcf-kit - append sample-level filters |
| hard-filtered | Any variant site flagged by the soft filters is removed, Any sample genotype with a soft-filter flag are set to missing | processed variant set | Same as soft-filter | CeNDR | BCFtools - remove soft-filter sites |
| PopGen | Processed variant set for population genomics analyses | pi;<br>tajima's D;<br>Fst;<br>phylogeny;<br>neighbor-net;<br>admixture;<br>treemix;<br>haplotype | No missing genotypes sites;<br>keep one variant in high LD blocks (LD > 0.95) | **Supplemental Data 4** | BCFtools - remove sites with missing genotypes;<br>plink - LD pruning |

**Supplemental Table 2 - Description of variant sets used in this study.** The additional filters applied to the PopGen VCF for specific uses are described in methods.

| Field | Description | Type (units; if applicable) | Example values |
|---|---|---|---|
| **C-label** | A label associated with a collection and obtained by scanning the barcode | Text | C-0001 |
| **Sample photo** | A photograph of the sample environment | Photograph | See figure 3C |
| **Substrate** | The sample substrate as determined in the field | Categorical | Leaf Litter, Fungus, orFlower |
| **Substrate notes** | Additional notes regarding a collected substrate | Text | Turned over log |
| **Landscape** | The type of environment from which a sample was obtained | Categorical | Wild forest<br>Wild grassland |
| **Sky View** | The visibility of the sky from the perspective of the sample collected | Categorical | Full<br>Partially Obstructed<br>Obstructed |
| **Gridsect (optional)** | Whether the sample was part of a gridsect | Yes / No | Yes<br>No |
| **Gridsect direction (optional)** | Only applies to gridset samples - Defines one of six directions collected within a gridsect | Categorical (degrees) | A, B, C, D, E, F |
| **Gridsect radius (optional)** | Only applies to gridsect samples - Defines the distance a sample was collected from the center | Categorical (meters) | 0 - Center<br>1<br>2<br>3 - Outer Circle |
| **Substrate temperature** | The temperature of the substrate as determined by local measurement | Numeric (ºC) | 16.4 |
| **Substrate moisture** | Moisture as determined by local measurement | Numeric (%) | 25 |
| **Ambient temperature** | The temperature of the environment | Numeric (ºC) | 18.2 |
| **Ambient humidity** | The humidity of the environment | Numeric (%) | 50 |

**Supplemental Table 3 - Nematode field sampling data form.** Sampling data are entered into fields of the data form and stored in a cloud database at Fulcrum ® (http://www.fulcrumapp.com).

| Field | Description | Type (units; if applicable) | Example values |
|---|---|---|---|
| **C-label** | The sample-collection bag | Text | C-0001 |
| **Worms on sample** | Whether worms were found associated with a sample | Yes / No | Yes |
| **Approximate number of worms (optional)** | An estimate of the number of worms found associated with a sample | Categorical | Very Few (1-3)<br>Few (4-10)<br>Some (11-25)<br>Proliferating (25+) |
| **Dauers on Sample** | Dauers were observed associated with sample | Yes / No | Yes<br>No |
| **Males Observed** | Males observed associated with sample | Yes / No | Yes<br>No |
| **S-labeled Plates** | A list of S-labeled plates; Each S-label corresponds to a single worm isolate | List | S-00001<br>S-00002 |

**Supplemental Table 4 - Nematode isolation data form.** Nematode isolation data are entered into fields of the data form. All data associated with the substrate, C-label, and S-label are linked and stored in a cloud database at Fulcrum®.

| Pipeline | Description | Availability |
|---|---|---|
| trimmomatic-nf | Performs trimming to remove poor quality sequences and technical sequences such as adapters. It should be used with high-coverage genomic DNA | http://github.com/andersenlab/trimmomatic-nf |
| alignment-nf | performs alignment for wild isolate sequence data on strain and isotype levels, and output BAMs and related information | https://github.com/AndersenLab/alignment-nf |
| concordance-nf | The concordance pipeline is used to detect sample swaps, identify samples with quality issues, and determine which wild isolate strains should be grouped together as an isotype | https://github.com/AndersenLab/concordance-nf |
| wi-nf | Calls variants for wild *C. elegans* isolates | https://github.com/AndersenLab/wi-nf |

113 **Supplemental Table 5 - Nextflow pipelines used in our study.**
