## Supplemental File 1 for "Deep sampling of Hawaiian *Caenorhabditis elegans* reveals high genetic diversity and admixture with global populations"

**Supplemental File 1 - Declaration of a new *Caenorhabditis* species**

The electronic edition of this article conforms to the requirements of the amended International Code of Zoological Nomenclature (ICZN), and hence the new names contained herein are available under that Code from the electronic edition of this article. This published work and the nomenclatural acts it contains have been registered in ZooBank, the online registration system for the ICZN. The ZooBank LSIDs (Life Science Identifiers) can be resolved and the associated information viewed through any standard web browser by appending the LSID to the prefix ''http://zoobank.org/''. The LSID for this publication is: urn:lsid:zoobank.org:pub:DBB22717-DA3E-4B44-A381-DDD89B4CF1BD. The electronic edition of this work was published in a journal with an ISSN.

*Caenorhabditis oiwi* Crombie et al. sp. n.

We isolated and identified a new *Caenorhabditis* species that we named *Caenorhabditis oiwi* sp. n. for the Hawaiian word meaning native. Here, we justify the species status of *C. oiwi* sp. n. based on molecular barcodes and biological species inference from mating experiments. The type isolate for *C. oiwi* sp. n. is strain ECA821. We also made an isogenized version of ECA821 by ten generations of sib mating (named ECA1100). The species reproduces sexually with males and females. The ITS2 sequence from ECA1100 *C. oiwi* sp. n. (Genbank Accession: MN056420) differs from that of all previously described *Caenorhabditis* species for which such information is available (Félix et al., 2014; Huang et al., 2014; Kiontke et al., 2011; Stevens et al., 2019). Note that these ribosomal DNA sequences might vary slightly within the species. Based on molecular data, *C. oiwi* sp. n. falls into the Elegans supergroup of *Caenorhabditis* (Kiontke et al., 2011) with the closest known species being *C. kamaaina* (Félix et al., 2014). Reciprocal mating experiments of *C. oiwi* sp. n. ECA821 with the *C. kamaaina* type isolate QG122 did not yield any viable progeny. *C. kamaaina* was previously described as a sister species to the *Japonica* group but was recently placed as the most basally diverging species in the Elegans group (Kiontke et al., 2011; Stevens et al., 2018). The discovery of *C. oiwi* sp. n. might help with resolving the shifting topology in this part of the *Caenorhabditis* phylogenetic tree.

The type isolate ECA821 was collected in August of 2017 from the Island of Oahu, Hawaii (21.33611°N, -157.7999°W) where it was isolated from a cluster of freshly fallen flowers. ECA821 is deposited as a cryo-preserved living stock at the *Caenorhabditis* Genetics Center. Isolate ECA821 is deposited in the NYU Rhabditid Collection and was used to study the morphology of the species (**Supplemental Figure 12; Supplemental Figure 13**). In agreement with the similarity of their rRNA sequences, *C. oiwi* sp. n. and *C. kamaaina* are at present morphologically indistinguishable. Both species show the common features of the Elegans group of *Caenorhabditis* (Sudhaus and Kiontke, 2007). Their lips are separate; the stoma is long and bears three flaps of moderate size at the metastegostom (**Supplemental Figure 12A-B**). The male tail shows the typical heart-shaped, anteriorly closed fan (bursa) with a serrated edge and a shallow terminal notch (**Supplemental Figure 13A, E**). The nine pairs of rays are arranged as is typical for the Elegans group with two pairs of rays positioned precloacally and the tips of ray pairs v1 are attached to the dorsal side of the fan. The anterior dorsal ray (ad) is in position five and the posterior dorsal ray (pd) in position seven. The tips of the sixth pair of rays (v5) are embedded in the cuticle. Rays v4 are much thinner and always shorter than ad, a character that distinguishes *C. oiwi* sp. n. and *C. kamaaina* from most species of the Elegans group (but not all; *C. doughertii, C. tropicalis* and *C. nigoni* also have a narrower and shorter ray v4). Several species of the Japonica group show modified rays v4. In *C. japonica, C. nouraguensis, C. panamaensis* and *C. waitukubuli,* rays v4 are much shorter than the ad rays. In *C. becei* and *C. macrosperma,* rays v4 are only slightly shorter than the ad rays, but not as skinny as in *C. kamaaina* and *C. oiwi.* The spicules are slender and their tip is pointed. The gubernaculum shows the usual forked distal tip and lateral ears (**Supplemental Figure 13C, D**), but both are more prominent than in most other species of the Elegans group. Here, only *C. inopinata* and *C. brenneri* have equally solid lateral ears and distal forked tip. The morphology of the females

(**Supplemental Figure 12B-F**) is in agreement with that of the stem species pattern of the Elegans group (Sudhaus and Kiontke, 2007).
